# Lymphatic endothelial cell-targeting lipid nanoparticles delivering VEGFC mRNA improve lymphatic function after injury

**DOI:** 10.1101/2024.07.31.605343

**Authors:** Eleftheria Michalaki, Rachel Chin, Kiyoung Jeong, Zhiming Qi, Lauren N. Liebman, Yarelis González-Vargas, Elisa Schrader Echeverri, Kalina Paunovska, Hiromi Muramatsu, Norbert Pardi, Beth Jiron Tamburini, Zoltan Jakus, James E. Dahlman, J. Brandon Dixon

## Abstract

Dysfunction of the lymphatic system following injury, disease, or cancer treatment can lead to lymphedema, a debilitating condition with no cure. Advances in targeted therapy have shown promise for treating diseases where conventional therapies have been ineffective and lymphatic vessels have recently emerged as a new therapeutic target. Lipid nanoparticles (LNPs) have emerged as a promising strategy for tissue specific delivery of nucleic acids. Currently, there are no approaches to target LNPs to lymphatic endothelial cells, although it is well established that intradermal (ID) injection of nanoparticles will drain to lymphatics with remarkable efficiency. To design an LNP that would effectively deliver mRNA to LEC after ID delivery, we screened a library of 150 LNPs loaded with a reporter mRNA, for both self-assembly and delivery *in vivo* to lymphatic endothelial cells (LECs). We identified and validated several LNP formulations optimized for high LEC uptake when administered ID and compared their efficacy for delivery of functional mRNA with that of free mRNA and mRNA delivered with a commercially available MC3-based LNP (Onpattro™). The lead LEC-specific LNP was then loaded with VEGFC mRNA to test the therapeutic advantage of the LEC-specific LNP (namely, LNP7) for treating a mouse tail lymphatic injury model. A single dose of VEGFC mRNA delivered via LNP7 resulted in enhanced LEC proliferation at the site of injury, and an increase in lymphatic function up to 14-days post-surgery. Our results suggest a therapeutic potential of VEGFC mRNA lymphatic-specific targeted delivery in alleviating lymphatic dysfunction observed during lymphatic injury and could provide a promising approach for targeted, transient lymphangiogenic therapy.

**One Sentence Summary:** Development of a novel lymphatic endothelial cell-targeting lipid nanoparticle via *in vivo* screening for mRNA delivery improves lymphatic regeneration and function after injury.

## INTRODUCTION

The lymphatic system helps maintain fluid balance in the body by draining excess interstitial fluid from tissues and depositing it into the bloodstream (*1*). Lymph flow is driven by both active lymphatic wall pumping and transient drops in fluid pressure due to contraction of surrounding tissue to promote uptake of fluid from interstitium to the initial lymphatic capillaries and through the lymphatic network (*2–4*). Specialized lymphatic muscle lines the collecting lymphatic vessels (LVs) downstream of initial LVs (*5*). Sections of LVs, where smooth muscle is present (lymphangions), perform spontaneous muscle contractions. The presence of lymphatic valves (*4, 6*), in conjunction with the active contraction cycle of individual lymphangions, makes the intrinsic pump highly effective for promoting unidirectional flow (*2*).

Damaged LVs become leaky and their efficiency for pumping fluid is reduced, leading to the excessive accumulation of fluid, macromolecules, and leukocytes (*7*). This accumulation, caused by lymphatic impairment, is associated with diseases including edema, fibrosis, obesity, cardiovascular disease, and neurological disorders (*7–10*). Furthermore, dysfunction of the lymphatic system can result in poor immune function and lead to inflammation (*11*), autoimmune diseases (*12*), impaired wound healing (*8*), and even tumor progression (13). Therefore, the lymphatic system is not only important for fluid transport, but also in modulating immune function (*12*). As an example, lymphatic endothelial cells (LECs), comprising lymphatic capillaries, influence dendritic cell maturation (*12, 14*).

Given the significant impact of the lymphatic system and LECs on multiple pathophysiological conditions, several studies have explored various treatments to address lymphatic dysfunction. Clinically, the amelioration of lymphatic dysfunction in disease conditions focuses on palliative (*15–18*) or surgical treatments, such as vascularized lymph node transplantation (*19–24*). However, patients receive these treatments for extended periods with high costs, and their efficacy remains limited, indicating the need for a pharmacotherapy strategy to achieve efficient therapy *in vivo* (*1*).

Reduction of inflammation has been used for treatment of lymphatic disfunction (*20*). For instance, ketoprofen, a non-steroidal anti-inflammatory drug (NSAID), has been shown to not only inhibit the inflammatory pathways of both cyclooxygenase (COX) and 5-lipoxygenase (5-LO) but also to promote lymphatic repair (*25*). However, NSAIDs carry the risk of heart attack or stroke (*26*). In addition of the inhibition of inflammatory pathways, growth factors, such as fibroblast growth factor 2 (FGF2), hepatocyte growth factor (HGF), retinoic acid (RA), and vascular endothelial growth factor C (VEGFC), have been studied in preclinical models as potential agents to induce lymphangiogenesis and improve lymphatic function. FGF2 and RA, however, induce lymphangiogenesis indirectly by upregulating VEGFC expression (*27*). VEGFC induces lymphangiogenesis by binding to vascular endothelial growth factor receptor 3 (VEGFR3) (*28, 29*). VEGFC-mediated lymphangiogenesis expands the lymphatic network, limits inflammation (*14*), and improves lymphatic function (*20*).

Utilizing nanomedicine for targeted drug delivery improves the effective dose at the target tissue while avoiding off-target effects. Specifically, nanoparticles (NPs) have been recently utilized as robust delivery agents by encapsulating or attaching therapeutic drugs and distributing them to target tissues (*30–32*). To deliver VEGFC to LECs, VEGFC protein-loaded biodegradable NPs have been recently studied with a variety of approaches, including poly lactic-co-glycolic acid nanosphere (*33*), gelatin hydrogel (*34*), and Mesenchymal stem cells (MSCs) derived-exosomes (*35*), and antibody conjugation to LNPs (*36*).

In addition, advances in genomics have led to the development of targeted gene therapies (*37*). To date, siRNA therapies have shown the most promise in patients with infectious and cardiometabolic diseases (*38, 39*), while adenovirus-based platforms only recently were utilized for patients with lymphatic dysfunction – a clinical trial based on an adenovirus-based VEGFC delivery platform (Lymfactin®) entered Phase II (NCT03658967) (*40, 41*). Although siRNA- and adenovirus-based approaches are currently the most clinically advanced therapies, mRNA-based platforms offer a transient method of delivery compared to protein therapeutics (*42*) or adenovirus and adeno-associated virus-based platforms (*43, 44*). RNA delivery vehicles are designed to protect the nucleic acid and transport it to the target cell, and LNPs can be a powerful vehicle for relevant cargo delivery (*45*).

In conjunction with rapid NP synthesis, high-throughput *in vivo* NP screening methods allow scientists to track many LNPs simultaneously. Specifically, high-throughput DNA barcoding systems have been developed that allow analysis of >100 LNPs *in vivo* (*46–50*). Notably, simultaneous administration of many LNP formulations overcomes challenges associated with expensive *in vivo* screening, and existing physical (e.g., brain accessibility, LNP disassembly) and physiological (e.g., undesired LNP binding to serum proteins) barriers (*47*). Several barcoding assays have been reported, a subset of which quantify functional mRNA delivery (i.e., delivered mRNA turning into protein) mediated by many LNPs at once. One such assay is called Species Agnostic Nanoparticle Delivery Screening (SANDS) (*48*), which measures the functional delivery of mRNA encoding an anchored VHH antibody (aVHH) (*49*). Given the implication of the lymphatic system in various disease conditions and the challenges of lymphatic-specific targeting, the multiplexity of DNA barcoding technology combined with the versatility of LNPs may be a means to accelerate the development of new lymphatic-specific genetic therapies.

Here we used SANDS to study how 150 different LNP formulations delivered mRNA and identified several LNPs *in vivo* that deliver functional mRNA to LECs. We then used the leading LNP (named LNP7) to deliver VEGFC mRNA to LECs, thereby improving lymphatic regeneration and function following lymphatic injury.

## RESULTS

### Identification of lead LEC-specific LNP candidates

To guide the delivery of VEGFC mRNA to lymphatics, we first used SANDS to identify an LEC-targeting LNP. Over three experiments we screened a library of 150 chemically distinct LNPs by varying LNP ionizable lipid, cholesterol, alkyl-tailed PEG, and helper lipids (e.g., DOPE, DSPC) (Fig. 1, Fig. S1). Each chemically distinct LNP was loaded with an mRNA encoding aVHH and a unique DNA barcode. After synthesis, each LNP formulation was evaluated *in vitro* for a series of criteria outlined in detail in the methods. Ninety-nine of these unique LNP formulation passed this stage and were selected for *in vivo* delivery. At each screening, LNPs were intradermally injected in each paw of 5 mice total (3 mice with LNPs for screening and 2 mice with saline for control). Between 12-16 hours after LNP injection the downstream LNs were collected and digested before using fluorescence-activated cell sorting (FACS) to sort LECs with high aVHH expression (i.e., cells in which aVHH mRNA was functionally delivered). LN were chosen for screening of LEC targeting instead of collecting lymphatic vessels, due to the substantially larger number of LEC within the subcapsular sinus of the LN compared to the afferent lymphatics. Sorted cells were pooled across all mice and all lymph nodes then sequenced to identify the DNA barcodes present within the cells, thereby identifying LNPs colocalized with cells in which functional delivery occurred.

**Fig. 1.**
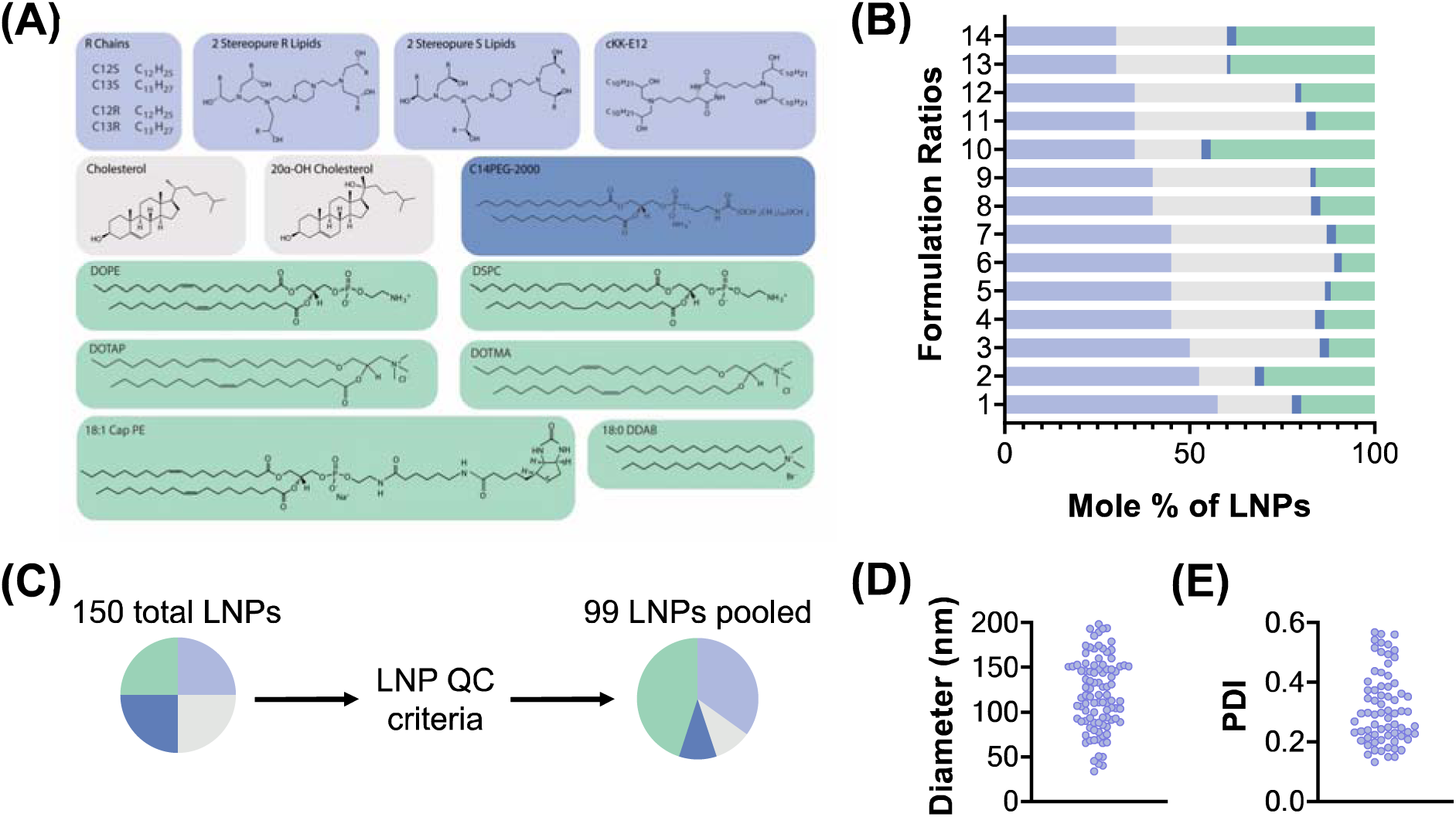
Screening different LNP formulations with FIND. **(A)** Main lipid components of the LNP libraries tested. **(B)** Each of the compounds was formulated using 14 molar ratios. **(C)** Of the 150 LNPs that were formulated, 99 passed the quality control (QC) criteria, with a diameter less than 200 nm as well as a stable autocorrelation curve. **(D)** Hydrodynamic diameters and **(E)** Polydispersity indexes (PDI) of all administered LNPs; the diameter of the LNP pooled control is within the range of the LNPs composing the pool.

After identification of LNPs that efficiently delivered functional aVHH mRNA to LECs residing in LNs, we identified six lead LNP candidates, which we named LNP1, LNP2, LNP3, LNP4, LNP7, and LNP11 (Fig. 2A, Table S1, and Table S2). We then confirmed the activity of the LNPs individually. We injected them into mice and quantified aVHH^+^ LECs in the axillary LN (ALN), brachial LN (BLN), and popliteal LN (PLN) using flow cytometry. We found that LNP1, LNP2, and LNP7 led to consistently high percentages of aVHH^+^ cells LECs in all examined LNs (ALN: LNP1 45%, LNP2 47%, LNP7 28% (Fig. 2B); BLN: LNP1 44%, LNP2 34%, LNP7 32% (Fig. 2C); PLN: LNP1 39%, LNP2 42%, LNP7 53% (Fig 2D)). LNP7 yielded the highest uptake by LECs in PLNs, which is the primary draining lymph node from the hindlimb and was used for subsequent experiments (Fig 2D).

**Fig. 2.**
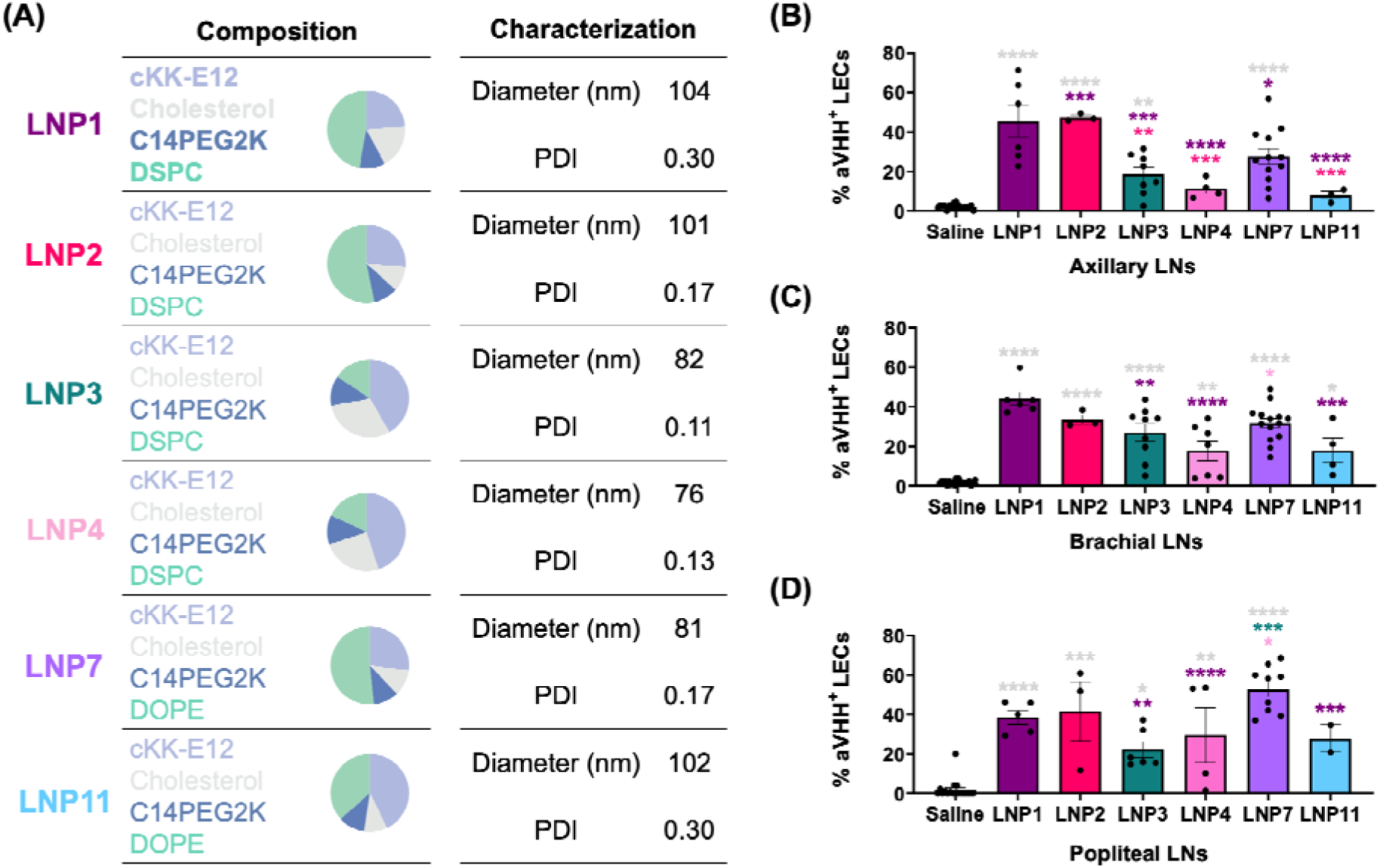
Validation of lead LEC-specific LNP candidates. **(A)** Formulation compounds, composition, hydrodynamic diameter (nm), and PDI of lead LEC-specific LNPs. **(B, C, D)** Percentage of aVHH^+^ LECs (after gating for Live/Dead, CD31^+^/PDPN^+^) from **(B)** ALN, **(C)** BLN, and **(D)** PLN after administration with saline (gray), LNP1 (dark purple), LNP2 (magenta), LNP3 (green), LNP4 (pink), LNP7 (purple), or LNP11 (blue). Each data point corresponds to an independent experiment (ALN: *N_Saline_* = 20, *N_LNP1_* = 6, *N_LNP2_* = 3, *N_LNP3_* = 8, *N_LNP4_* = 4, *N_LNP7_* = 13, and *N_LNP11_* = 3; BLN: *N_Saline_* = 22, *N_LNP1_* = 6, *N_LNP2_* = 3, *N_LNP3_* = 9, *N_LNP4_* = 8, *N_LNP7_* = 14, and *N_LNP11_* = 4; PLN: *N_Saline_* = 18, *N_LNP1_* = 5, *N_LNP2_* = 3, *N_LNP3_* = 6, *N_LNP4_*= 4, *N_LNP7_* = 9, and *N_LNP11_* = 2), and error bars represent the corresponding standard error of the mean (SEM). Color-coordinated asterisks above plots indicate a pairwise comparison for significance using a one-way ANOVA and Robust regression and Outlier removal (ROUT) method to identify and remove outliers, followed by a post-hoc test to correct for multiple comparisons with *p* < 0.05 (*), *p* < 0.01 (**), *p* < 0.001 (***), and *p* < 0.0001 (****).

### LNP7 provides superior functional mRNA delivery to LEC in both the draining lymphatic vessel and the draining lymph node

Next, we sought to demonstrate the superiority of LNP7, which led to the highest LEC-specific uptake in PLN, in delivering mRNA cargo to LECs *in vivo* in both the LEC of the afferent collecting lymphatic vessel and the draining lymph node. We compared LNP7 delivery of aVHH mRNA to LEC of the PLN to administration of saline, free mRNA (aVHH mRNA), and MC3-based LNPs (MC3) loaded with aVHH mRNA, an FDA-approved hepatocyte-targeting LNP formulation in Onpattro™ (*51, 52*) (Fig. 3A). In this study, we only injected one hindlimb, which allowed us to use the lymph nodes that drain the remaining limbs as controls for off-target delivery. After 12-16 hrs of administration, we isolated the PLN draining the injection site (PLN Injected) from the injection site and the ALN, the BLN, and the contralateral PLN, PLN collected (PLN Contralateral) from the non-injected sites and quantified the percentage of LEC that express aVHH using flow cytometry (Fig. 3, Table S3 and Table S4). LNP7 led to the highest mRNA delivery to LECs in the PLN Injected (saline 2%, Free aVHH 2%, MC3 5%, LNP7 31%) (Fig 3B). There was no significant functional mRNA delivery in vessels and nodes that drain the non-injected sites (ALN, BLN, and PLN Contralateral), demonstrating that the enhanced LEC delivery of LNP7 is specific to LEC of the lymph node that directly drain the injection site (Fig 3B). In an additional study, we dissected the afferent popliteal LVs (PLV) and upon quantification with FACS found that LNP7 consistently led to the highest cargo delivery to LECs in collecting LVs (saline 1%, Free aVHH 2%, MC3 10%, LNP7 37%) (Fig. 3C, Table S3). In addition, we investigated the delivery of LNPs in other endothelial and immune cell populations that reside in the draining and non-draining LNs using a 12-channel flow cytometry with a gating strategy detailed in the provided supplemental materials. aVHH expression remained low (below ∼15%) for all examined cell populations except for dendritic cells (cDC2 and cDC1), where we saw a significant uptake of LNP7 in the injected PLN (∼42% and ∼28%, respectively; Fig. S2f and g). aVHH expression also remained low for all cell populations in the non-draining lymph nodes (Fig. S2). We then evaluated LNP7 tolerability and found no evidence of overt toxicity LNP7 (Fig. S3).

**Fig. 3.**
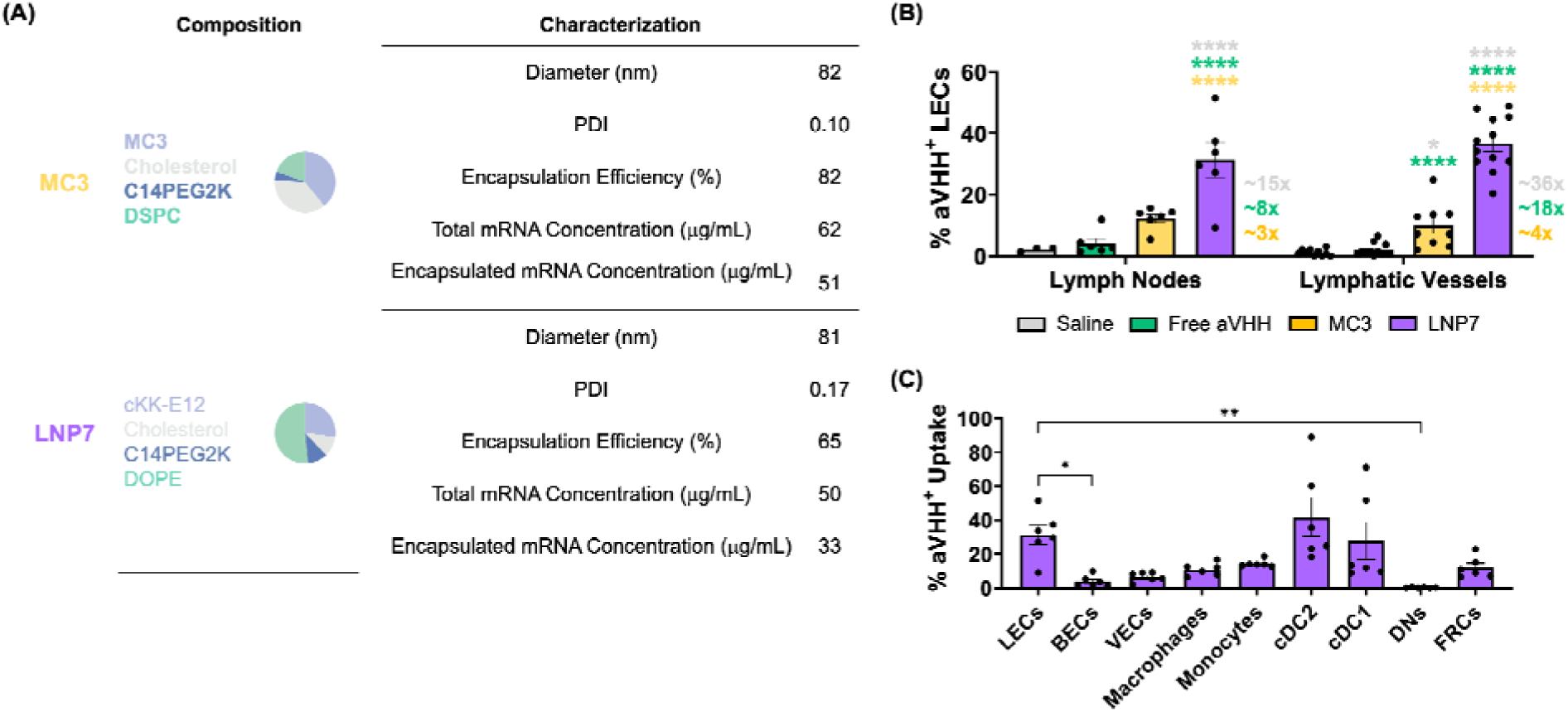
LNP7 provides superior delivery of mRNA to LEC and DCs within the draining lymph node and to LEC within the collecting lymphatic vessels. **(A)** Formulation compounds, composition, hydrodynamic diameter (nm), PDI, encapsulation efficiency (%), total mRNA concentration (μg/mL), encapsulated mRNA concentration (μg/mL) of MC3 and LNP7. **(B)** Percentage of aVHH^+^ LECs from PLN and PLV after delivery with saline (gray), free aVHH (green), MC3 (gold) and LNP7 (purple). **(C)** Percentage of aVHH^+^ cells upon delivery with LNP7 by different cell types within the draining lymph node, namely LECs, blood endothelial cells (BECs), vascular endothelial cells (VECs), macrophages, monocytes, dendritic cells (cDC2, cDC1), double negative cells (DNs), and fibroblastic reticular cells (FRCs). Each data point corresponds to an independent experiment (Lymph Nodes: *N_Saline_* = 3, *N_Free aVHH_* = 6, *N_MC3_* = 6, and *N_LNP7_* = 6; Lymphatic Vessels: *N_Saline_* = 10, *N_Free aVHH_*= 12, *N_MC3_* = 9, and *N_LNP7_* = 12), and error bars represent the SEM. **(B)** Color-coordinated asterisks above plots indicate a pairwise comparison for significance using a two-way ANOVA with Tukey’s multiple comparisons test with *p* < 0.05 (*) and *p* < 0.0001 (****). **(C)** Solid lines above plots indicate a pairwise comparison for significance using a one-way ANOVA with Tukey’s multiple comparisons test with *p* < 0.05 (*) and *p* < 0.01 (**). Identification of the respective cell populations (after Live/Dead gating) was as follows: LECs: CD45^-^/CD31^+^/PDPN^+^, VECs: CD45^-^/CD31^+^/PDPN^-^/CD54^+^, BECs: CD45^-^/CD31^+^/PDPN^-^/CD309^+^, FRCs: CD45^-^/CD31^-^/PDPN^+^, DNs: CD45^-^/CD31^-^/PDPN^-^, Monocytes: CD45^+^/CD11b^+^/CD64^+^/F4-80^-^, Macrophages: CD45^+^/CD11b^+^/CD64^-^/F4-80^+^, cDC2: CD45^+^/CD11c^+^/MHCII^+^/CD11b^+^, and cDC1: CD45^+^/CD11c^+^/MHCII^+^/CD11b^-^.

### Optimization of LNP VEGFC mRNA dose for enhancing lymphatic repair and function

After identifying several LNP formulations with high lymphatic delivery when administered intradermally compared to free mRNA and MC3-based LNPs, we next investigated whether LNP7 would improve the potential therapeutic efficacy of VEGFC mRNA delivery for lymphatic regeneration and restoration of lymphatic pump function. We hypothesized that local VEGF-C mRNA delivery using LNP7 to LEC within vessels draining the site of injury would improve therapeutic outcomes compared to no treatment, free VEGF-C mRNA, or delivery with VEGF-C mRNA with an LNP with poor LEC delivery. To test this, we decided to utilize a mouse tail lymphatic injury model previously developed by our lab, where one chain of lymphangions is damaged, while the parallel lymphangion chain on the adjacent side of the tail remains intact.

First, we determined the potential effect of varying doses of LNP7-loaded VEGFC mRNA delivery on lymphatic function and regeneration after lymphatic injury. We used 4 different dosages of VEGFC mRNA (0.04 μg, 0.2 μg, 1 μg, and 5 μg) loaded into LNP7 and LNP7 without VEGFC mRNA (Empty LNP - LNP7 loaded with aVHH mRNA instead of VEGF-C mRNA), to serve as a control. Mice were administered a single injection of LNP7 at the respective dose 3 days after injury.

To analyze how lymphatic transport in the intact collecting vessel changed over time after treatment, we utilized NIR imaging to quantify functional metrics of lymphatic contractility both before injury and 7 days after surgery in each treatment group. Administration of 5 μg significantly increased the packet frequency of lymphatic contraction, while lower VEGFC mRNA doses (namely, 0.04 μg, 0.2 μg, and 1 μg) had no significant effect on lymphatic function (Fig. 4).

**Fig. 4.**
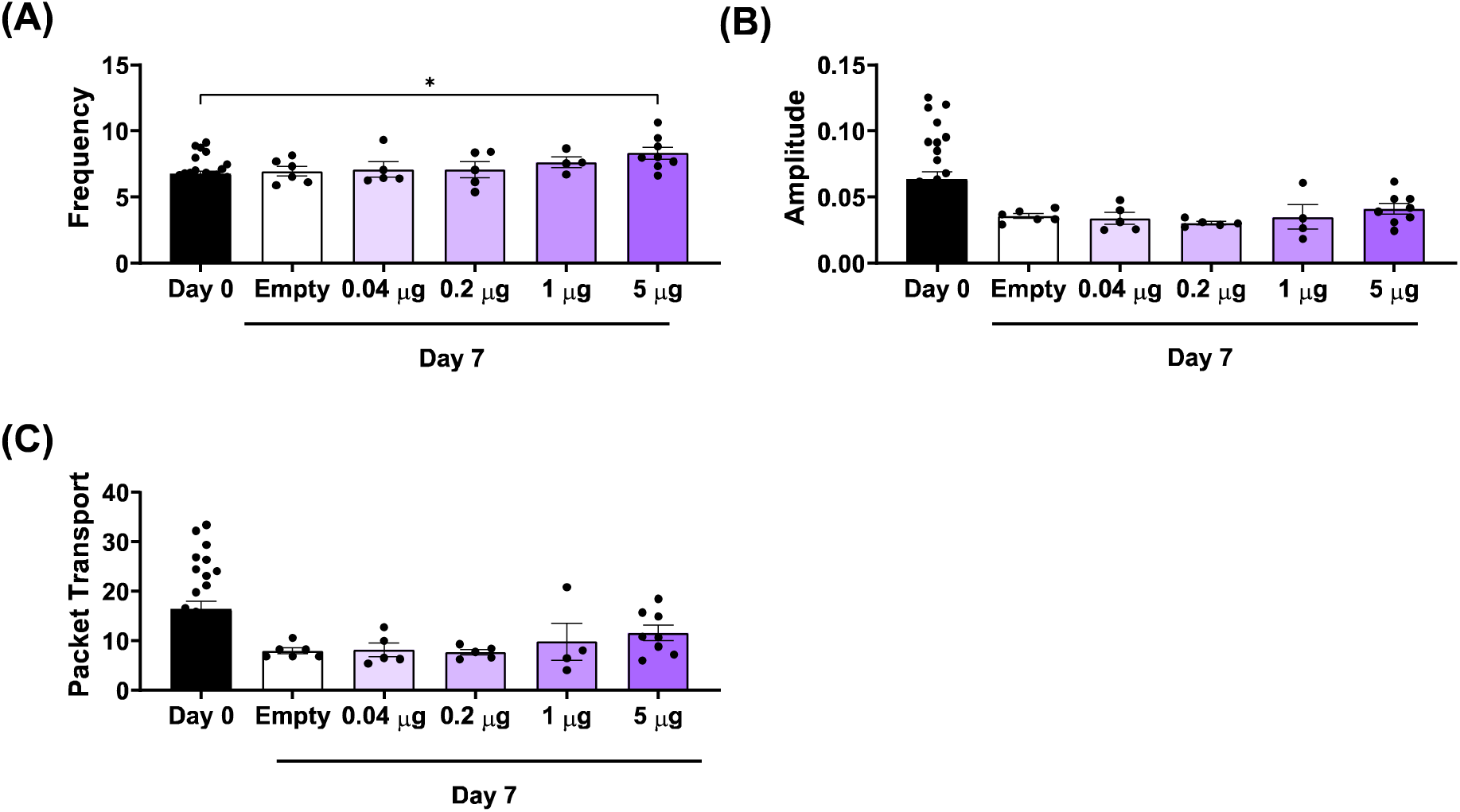
LNP7 loaded with 5 μg VEGFC mRNA significantly increased LV pumping frequency 7 days after mouse tail lymphatic injury *in vivo*. **(A)** Frequency, **(B)** amplitude, and **(C)** packet transport for empty (black), 0.04 μg (light purple), 0.2 μg (purple), 1 μg (dark purple), and 5 μg (darker purple) mRNA-LNP7 treatment measured 7 days post-surgery. Each data point corresponds to an independent experiment (*N_Empty_* = 6, *N_0.04_* _μ_*_g_* = 5, *N_0.2_* _μ_*_g_* = 5, *N_1_* _μ_*_g_* = 4, and *N_5_* _μ_*_g_* = 8), and error bars indicate the corresponding SEM. Solid lines above the plots indicate a pairwise comparison for significance using one-way ANOVA with Tukey’s multiple comparisons test with *p* < 0.05 (*), *p* < 0.01 (**), and *p* < 0.001 (***).

We next evaluated lymphangiogenesis by investigating histological changes in the tail. Circular cross-sections were taken from the wound site and the distal portion of the tail from blank and 5 μg treatment groups and stained for podoplanin (PDPN), an LEC marker, and EdU to measure cell proliferation. We demonstrated the presence of PDPN-positive and EdU-positive cells in both the wound and distal sites in the Empty and 5 μg groups (Fig. 5A). Upon quantification, we observed higher PDPN/EdU colocalization only in the wound site of mice treated with 5 μg VEGFC mRNA-LNP7 (Fig. 5B).

**Fig. 5.**
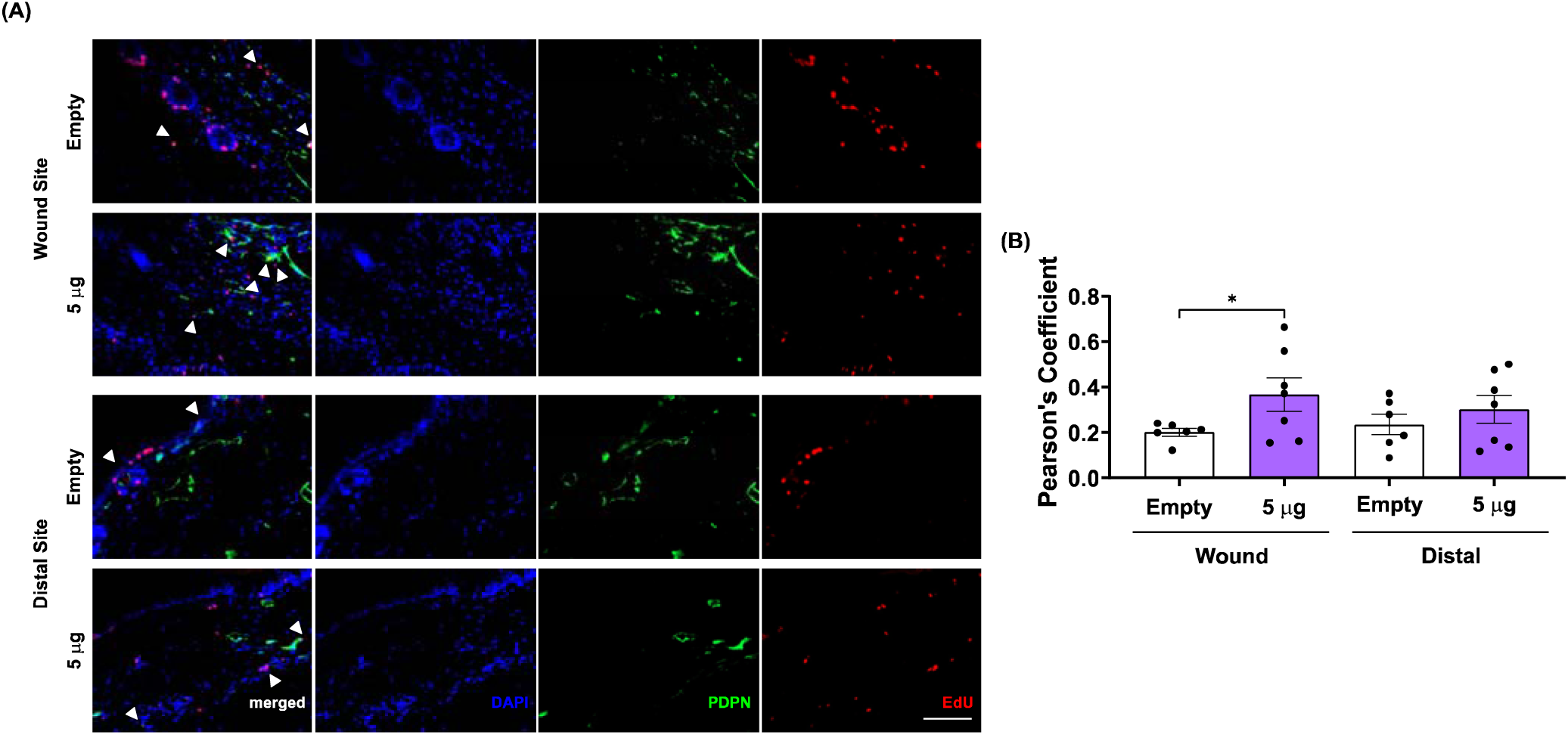
VEGFC mRNA overexpression significantly increases PDPN/EdU colocalization at the wound site 7 days after lymphatic injury. **(A)** Representative images of merged, DAPI (blue), PDPN (green), and EdU (red) in tail sections for Empty and 5 μg treatment group at the wound and distal sites 7 days post-surgery. Arrows indicate EdU and PDPN double positive LEC (20x objective; Scale bar = 100 μm). Contrast was enhanced post-acquisition for ease of viewing and was performed identically across all images. **(B)** Pearson’s coefficient in Empty (black) and 5 μg (purple) groups 7 days post-surgery in the wound and distal sites. Each data point corresponds to the average multiple sections (2-3 sections /mouse) taken from each independent experiment (*N_Empty_*= 6 mice and *N_5_* _μ_*_g_* = 7 mice), and error bars indicate the corresponding SEM. Solid lines above plots indicate a pairwise comparison for significance using a nested one-way ANOVA with Tukey’s multiple comparisons test with *p* < 0.05 (*).

### LNP7 delivery of VEGF-C mRNA improves lymphatic function up to 14 days after injury

After demonstrating that a VEGFC mRNA dosage of 5 μg increases lymphatic pump function and lymphangiogenesis 4 days after administration, we administered this same dose to determine the persistence of improvement on lymphatic pump function and the benefit compared to an LNP that does not target LECs. Mice received either empty LNP7 (Empty), 5 μg MC3-loaded with VEGFC mRNA (VEGFC mRNA-MC3), and 5 μg LNP7-loaded with VEGFC mRNA (VEGFC mRNA-LNP7).

Lymphatic transport metrics were measured pre-surgery and 7 and 14 days post-surgery. Although both VEGFC mRNA-MC3 and VEGFC mRNA-LNP7 therapies improved lymphatic contractile activity compared to Empty treatment 7 days post-injury, only VEGFC mRNA-LNP7 maintained improvement at 14 days post-injury (Fig. 6A). Thus, VEGFC mRNA-LNP7 treatment increases lymphatic function by increasing LV contraction frequency compared to the Empty and VEGFC mRNA-MC3 treatments 14 days post-injury (Fig. 6A-D).

**Fig. 6.**
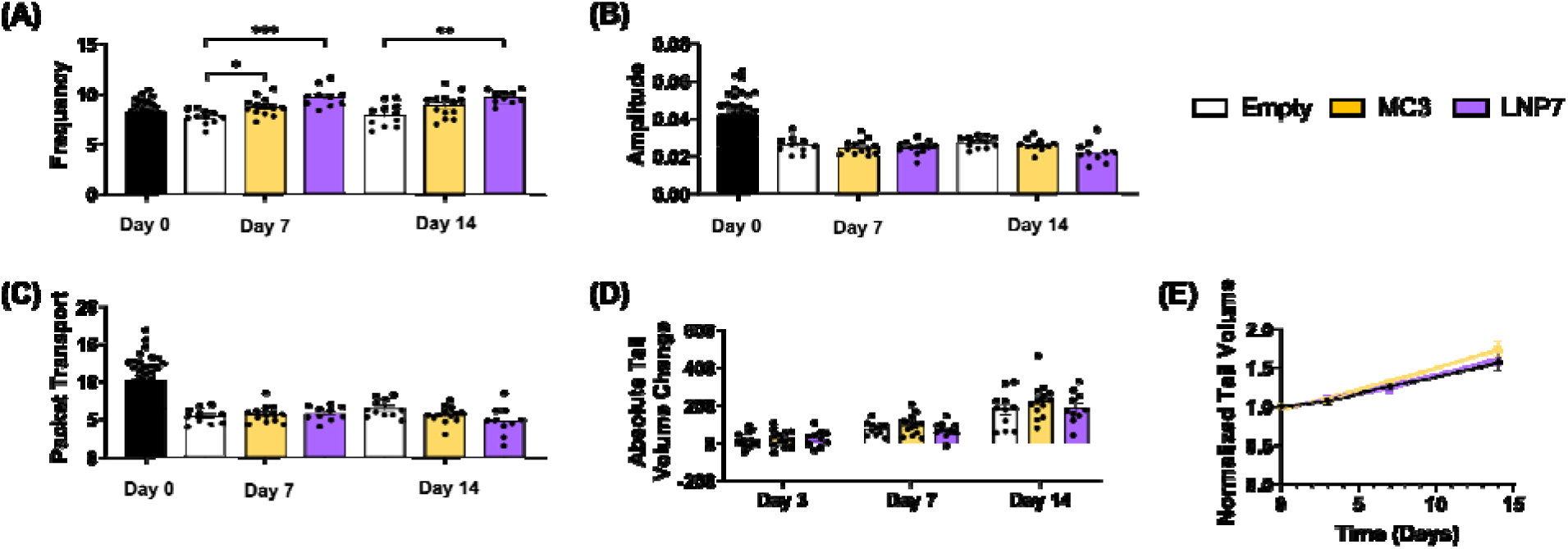
LNP7 loaded with VEGFC mRNA significantly increased LV pumping frequency 14 days after mouse tail single LV ligation surgery *in vivo*. **(A)** Frequency, **(B)** amplitude, **(C)** packet transport, **(D)** absolute tail volume change, and **(E)** normalized tail volume for empty (black), VEGFC mRNA-MC3 (5 μg) (gold), and VEGFC mRNA-LNP7 (5 μg) (purple) 14 days post-surgery. A linear regression model was fit to the data to find the best-fit value of the slope and intercept (y = β_1_x + β_0_: y*_Empty_* = 0.04183*x + 0.9722, y*_MC3/VEGFC_* = 0.05497*x + 0.9562, y*_LNP7/VEGFC_*= 0.04563*x + 0.9662). Each data point corresponds to an independent experiment (*N_Empty_* = 11, *N_MC3/VEGFC_* = 13, and *N_LNP7/VEGFC_* = 10), and error bars indicate the corresponding SEM. Solid lines above the plots indicate a pairwise comparison for significance using Mixed-effects analysis with Tukey’s multiple comparisons test and robust regression and outlier removal (ROUT) method to identify and remove outliers with *p* < 0.05 (*), *p* < 0.01 (**), and *p* < 0.001 (***).

Next, we examined the effect of the treatments on tail swelling. To do so, we used the tail images obtained 3, 7, and 14 days post-surgery. Modeling the tail as a series of truncated cones (*53*), we calculated total tail volume and determined the corresponding percentage of tail volume change. None of the treatments led to statistically significant changes in tail swelling (Fig. 6E). In addition, after fitting a linear regression model (y = β_1_x + β_0_), we found no difference in the swelling rate of mice receiving Empty, VEGFC mRNA-MC3, or VEGFC mRNA-LNP7 treatments (β_1, Empty_= 0.04183, β_1, MC3/VEGFC_ = 0.05497, and β_1, LNP7/VEGFC_ = 0.04563) (Fig. 6F).

Therefore, the use of VEGFC mRNA-LNP7 is not sufficient to reduce swelling over this time course in this particular animal model. This aligns with our previous reports and is likely due to the presence of an intact outflow route from the tail minimizing the impact of the surgery on swelling (54).

Given the benefit of VEGFC treatment on lymphatic function, we next sought to determine if there was any effect on lymphangiogenesis by investigating histological changes of LEC in the tail. Cross-sections were isolated from the wound site and the distal portion of the tail from each treatment group and stained for PDPN and EdU. As in the dose optimization study, we demonstrated the presence of PDPN-positive LECs in both the wound and distal sites in the three treatment groups (Fig. 7A and Fig. S4). Neither VEGFC mRNA-MC3 nor VEGFC mRNA-LNP7 treatment significantly modified the lymphatic network density 14 days after mouse tail single LV surgery in the wound or distal sites *in vivo* (Fig. 7 B-D).

**Fig. 7.**
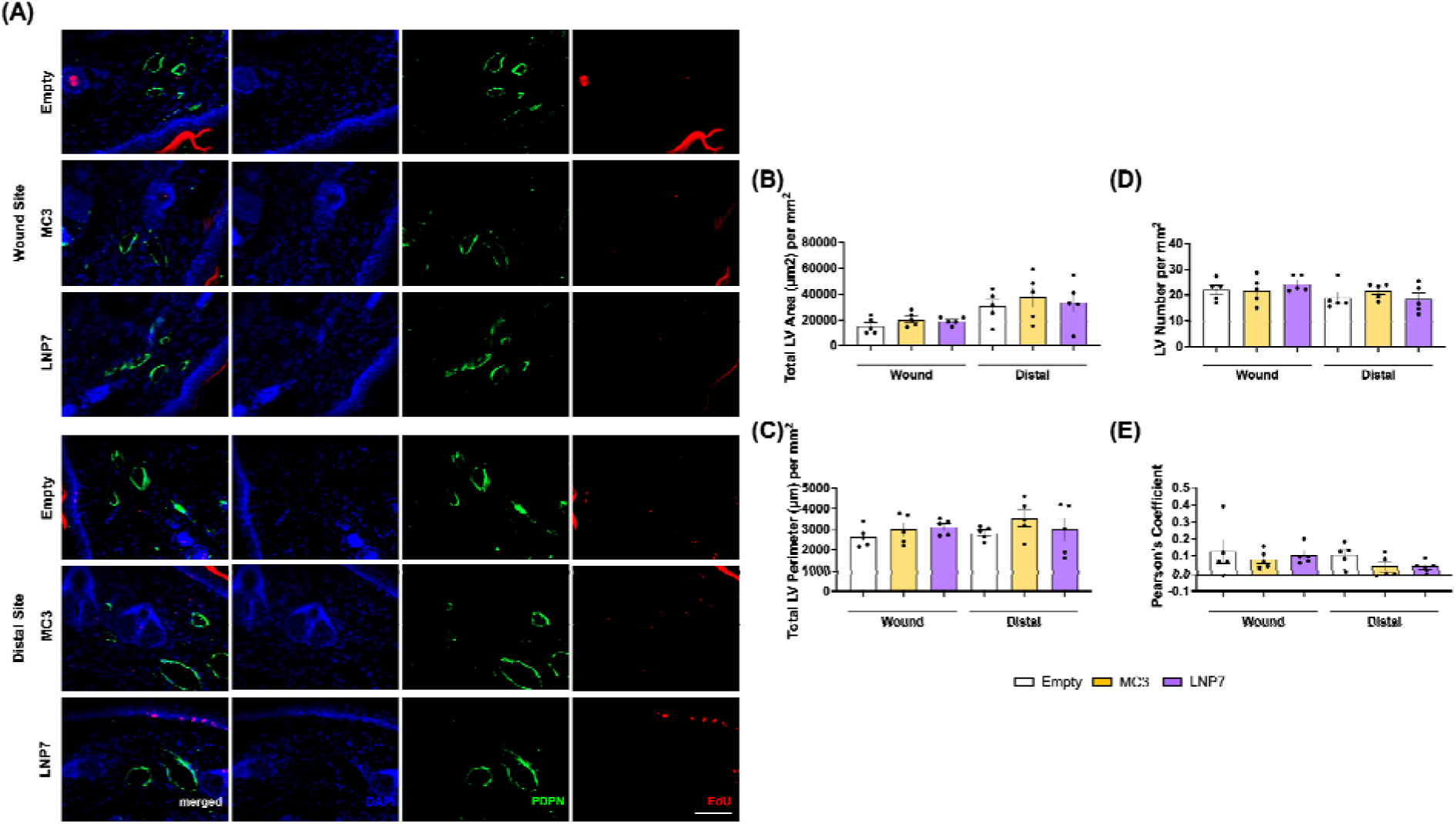
VEGFC mRNA delivery does not affect LV density or PDPN/EdU colocalization 14 days after mouse tail single LV ligation surgery *in vivo*. (**A**) Representative images of merged, DAPI (blue), PDPN (green), and EdU (red) in LVs for empty, VEGFC mRNA-MC3 (5 μg), and VEGFC mRNA-LNP7 (5 μg) in wound and distal sites 14 days post-surgery. (20x objective; Scale bar = 100 μm.) Contrast was enhanced post-acquisition for ease of viewing. Quantification of the **(B)** total LV area, **(C)** total LV perimeter, **(D)** total LV number per mm^2^, and **(E)** Pearson’s coefficient for empty (black), VEGFC mRNA- MC3 (5 μg) (gold), and VEGFC mRNA-LNP7 (5 μg) (purple) 14-days post-surgery in the wound and distal sites. Each data point corresponds to the average of each independent experiment (*N_Empty_* = 5, *N_MC3/VEGFC_* = 5, and *N_LNP7/VEGFC_* = 5), and error bars indicate the corresponding SEM. No significant differences were detected by pairwise comparison for significance using a nested one-way ANOVA with Tukey’s multiple comparisons test with *p* < 0.05.

## DISCUSSION

The lymphatic system is part of the circulatory system (*1*), regulating tissue fluid balance and uptake. Failure to establish adequate tissue drainage results in lymphedema (*55, 56*), a condition for which there are currently no curative therapies (*57*). Despite the role of the lymphatic system in many pathologies, enhancing lymphatic drainage *in vivo* through targeted therapy has received little attention.

There have been increased efforts to utilize the lymphatic system as therapeutic modality for various pathological conditions (*9*). The combinatorial examination of the molecular mechanisms that govern lymphangiogenesis and key factors that dictate lymphatic function such as lymphatic drainage and pumping, could be applied towards efficient therapies. Here, we proposed the use of an innovative new technology for *in vivo* gene editing targeted at LECs to (i) screen and identify LNPs that target LECs, (ii) deliver functional mRNA into lymphatics, and (iii) utilize functional mRNA delivery for targeted therapy administration towards lymphatic regeneration in lymphedema.

Systemic endocrine therapies have caused vascular regression in endocrine organs and other tissues, necessitating use of more targeted solutions (*58–60*). Nanomedicine for functional mRNA delivery has emerged as a promising avenue to increase cell- and tissue-specific delivery. In lymphedema, although VEGFC administration is the most widely used and thoroughly investigated therapeutic for lymphatic-associated pathologies (e.g., BioBridge™ (61), Lymfactin® (*40, 41*)), its efficacy in treating the disease has produced mixed results (*19*). Thus, it seems that the development of tools to specifically target the lymphatic system may provide alternative therapeutic approaches towards the development of an effective lymphedema treatment.

These data support the hypothesis that LEC-specific LNPs delivering mRNA to LECs may be a useful therapeutic modality. Given the long-established role of VEGFC in lymphangiogenesis (*62,63*), LNPs carrying VEGFC mRNA to lymphatics suggests a promising and novel avenue for therapeutic purposes. VEGFC mRNA delivery in LECs using LNPs will lead to the transient enhancement of VEGFC (*53*); this could avoid problems associated with the long-term upregulation of VEGFC observed with other therapeutics. For example, permanent upregulation of VEGFC has been shown to promote cancer cell metastasis (*64*) and is associated with a leaky and dysfunction lymphatic vasculature. In contrast, LNP7 delivery of VEGFC mRNA led to enhanced proliferation of LEC at the site of injury downstream of LNP delivery, and was transient, occurring at 4 days after LNP treatment, but not persisting at 11 days following a single dose. While the study here only investigated the effect of a single dose, LNPs have been redosed every three weeks in patients for several years (*65*), suggesting that an optimized repeat LNP dosing schedule could be designed for different therapeutic contexts.

VEGFC is an ideal therapeutic cargo to showcase the utility of LEC targeting LNP therapy because of its role in regulating lymphangiogenesis (*62,63*) and that it has established effects on lymphedema (*66*). Also, the custom-made VEGFC mRNA used for our experiments has been previously been demonstrated to induce lymphatic growth and formation of a functional lymphatic network, restoring lymphatic function without adverse events in a mouse model of lymphedema (*67*).

Here, the versatility of SANDS facilitated identification of LEC-specific LNPs for functional targeted mRNA delivery. Our ability to identify LNPs that target LECs that reside in LNs and line collecting LVs may revolutionize targeted therapy in multiple pathological conditions. While large library screens were performed with LN LECs due to the much larger numbers of LEC that line the subcapsular sinus than that of the collecting vessels in the mouse, validation of the lead candidate LNP7 in collecting LECs provides evidence than properties favorable for uptake by LN LEC are similar to those in collecting vessels. Combining LNP screening with scRNA-seq (*68*) could be an interesting future endeavor to evaluate whether properties favorable for LEC delivery differ by various LEC subsets (*69*). The simultaneous high uptake of LNP7 by dendritic cells (cDC2 and cDC1), which have been shown to regulate immune responses during lymphedema progression (*70*), provide another cellular target that can be achieved with LNP7 mRNA delivery. In the context of this study, delivery of VEGFC mRNA to these “off-target” cells did not appear to have any obvious unwanted therapeutic effects. In fact, it is unclear the extent that LEC delivered vs. DC delivered mRNA deliver of VEGFC mRNA is ultimately responsible for the functional benefit of LNP7.

Additionally, the ability of LNP7 to carry mRNA cargoes of different sizes (200 kDa [aVHH mRNA: 231 kDa] to 700 kDa [VEGFC mRNA: 703 kDa]) demonstrates the versatility of this delivery vehicle. Here, we proposed the use of VEGFC mRNA towards a therapeutic effect for lymphedema, but the options for potential cargos are limitless. For example, future work could combine delivery of mRNA for both VEGFC and its LEC specific receptor VEGFR3 (*71*). Another approach would be to load LNPs with both mRNA and small molecules, such as the immunosuppressive drug Cyclosporine A (CsA), which assists in controlling LEC proliferation and migration (*72*), for the restoration of lymphatic function and ultimately the efficient treatment of lymphedema. We also use LNPs loaded with fluorescent cargo for diagnostic purposes. There are endless possibilities, and we hope our work to be the first of many studies to follow.

One limitation of the study is that different lead LEC-specific LNPs seem to have a preferential uptake by different LNs; for example, LNP3 more efficiently targets LECs in BLNs compared to LECs in PLNs. Thus, LNP7 is not likely a “one-LNP-fits-all” delivery method. We anticipate that different LNPs would lead to the highest uptake in different areas of interest, thus designing and characterizing LNPs specifically for the area of interest would lead to the best results. Another limitation is the observed LNP batch variability in tissue targeting. Varied dialysis and centrifugation methods can alter LNP stability; batch to batch consistency is an ongoing area of research with the LNP field. This method may require the need for “fresh” LNPs synthesized the day of injection, as this constrain was built into the experimental design, and the extent that LNP7 would still remain functional after several days of storage, and what those ideal storage conditions would be, remain outside the scope of this current study.

In conclusion, the *in vivo* screen of LNPs to identify an LEC-specific LNP seeks to establish a novel technique for cargo delivery to lymphatics. The development of a versatile tool targeting LECs could revolutionize targeted therapy in a variety of disease processes associated with the lymphatic system. Targeted therapy utilizing an LEC-specific LNP is *to our knowledge the first attempt* towards efficient mRNA delivery in LECs and its corresponding use as a therapeutic modality. Specifically, our technique seeks to: (i) improve current targeting and bioavailability, (ii) reduce cost associated with existing techniques (e.g., antibody conjugation) (*73*), and (iii) facilitate the development of lymphatic-specific therapies. We developed a minimally invasive, LEC-specific, and efficient method to trigger lymphatic regeneration that might present a promising therapeutic modality towards multiple pathological conditions.

## MATERIALS AND METHODS

### Study Design

The aim of this study was to develop an mRNA-LNP platform optimized for targeting lymphatic endothelial cells (LECs) and investigate its potential to improve lymphatic function in a lymphedema mouse model through the administration of LNPs loaded with VEGFC mRNA. The experimental design involved screening formulated LNPs with varying compositions in normal mice without lymphedema-inducing single lymphatic vessel (LV) ligation surgery. These LNPs were characterized with their formulation parameters and lymphatic specificity, leading us to determine that LNP7 is the most potential candidate. Subsequently, lymphedema-induced mice with single LV ligation surgery were administered VEGFC mRNA-LNP7 intradermally. The study measured lymphangiogenesis and lymphatic pump function using immunofluorescence and near-infrared (NIR) imaging. Control groups were included to compare the lymphatic specificity or therapeutic efficiency of LNP7 with other formulations. Animals in this study were strain- and age-matched and randomly assigned to treatment groups and were administered LNPs intradermally. *In vitro* assessment of cell viability and metabolic rate in human LECs was conducted to evaluate the toxicity of LNP7. Sample sizes and replicates for the studies displayed in the figure captions were based on previously published work (*58–61,74*). In lymphedema studies, quantification of tail volume and histological parameters was performed in a blinded manner by investigators to assess therapeutic efficacy. Outliers were removed by robust regression and outlier removal (ROUT) in several experiments as described in the methods.

### Animal studies

All procedures were approved by the Georgia Institute of Technology IACUC Review Board. Female C57Bl/6 mice aged 7 to 12 weeks (Jackson Laboratory, Bar Harbor, ME) were used for all animal studies. The sexual disparity is due to the disproportional occurrence of lymphatic injury in females (*75,76*). Animal weight was recorded before and after all procedures. Studies were carried out at the Physiological Research Labs, Georgia Tech, Atlanta, GA.

### mRNA design and production

aVHH (*77*) and VEGFC mRNAs (*67,78*) were designed and produced based on previous studies. The aVHH plasmid was ordered from DNA geneblock and linearized with Not-I HF (New England Biolabs), then PCR purified using a PCR clean-up kit (Qiagen). Transcribed aVHH mRNA was capped with RNA and added with a poly-A tail following the mScript kit instructions. The purification of aVHH mRNA was performed using the RNeasy kit (Qiagen) and treated with Antarctic Phosphatase (New England Biolabs) for 1 hour.

For VEGFC mRNA production, a plasmid encoding codon-optimized mouse Vascular Endothelial Growth Factor C was linearized and then an *in vitro* transcription reaction was performed using T7 RNA polymerase (Megascript, Ambion). The plasmid encoded a 101-nucleotide-long poly (A) tail. N1-methylpseudouridine (m1Ψ)-5′-triphosphate (TriLink) instead of Uridine-5′-triphosphate (UTP) was incorporated into the VEGFC mRNA. VEGFC mRNA was capped by using Cleancap (TriLink) and cellulose-purified as described (*78*). All mRNAs were analyzed by agarose gel electrophoresis, measured for concentration, and stored frozen at −20 °C.

### Nanoparticle and mRNA dosing

Before injection, all LNPs loaded with aVHH or VEGFC mRNA were characterized as described in Supplementary Materials and Methods (Fig. S5). Animals were anesthetized using inhaled isoflurane (5% induction, 2-2.5% maintenance). In this study, all LNPs were injected intradermally based on initial screenings, which showed higher uptake by LECs compared to intravenous administration (Fig. S6). During LNP screening and characterization with normal mice without single LV ligation surgery, we injected LNPs in each paw of mice with a 1.5 mg/kg dosage intradermally. In each screening run, the number of mice were 3 and 2 for LNPs and saline respectively.

To determine the dose response of LNP7, lymphedema-induced mice by single LV surgery were injected intradermally into the tip of the tail with varying dosages of VEGFC mRNA (0.04 μg, 0.2 μg, 1 μg, and 5 μg) (*67*) on day 3 post-single LV ligation surgery. Control animals were intradermally injected to the tip of the tail with matching volumes of saline. The number of mice were 6, 5, 5, 4, and 8 for saline, 0.04 μg, 0.2 μg, 1 μg, and 5 μg VEGFC mRNA-LNP7 respectively.

To monitor the therapeutic effect in lymphatic function by VEGFC mRNA-LNP7, lymphatic injury-induced mice by single LV surgery were injected intradermally to the tip of the tail with empty LNP7 and 5 μg (mRNA dosage) of VEGFC mRNA-LNP7 on day 3 post-single LV ligation surgery. The number of mice were 6 and 7 for saline and 5 μg of VEGFC mRNA-LNP7 respectively. The difference in number between groups for the various lymphatic injury experiments was due to some mice having to be withdrawn from the study due to IACUC endpoint criterion from poor tissue healing in response to the surgery which occasionally occurs due to the artery accidentally being injured during cauterization of the tail wound. It Is also worth noting that mice were randomized after injury to determine which therapeutic treatment they would receive.

### Tissue collection

For LNP screening and characterization, tails, LNs, and LVs were isolated 12-16 hrs after nanomedicine administration (Fig. S7). During lymphedema studies, tails were collected 7- and 14-days post-injury for all treatment groups. For each tail, two 1-cm long tissue samples were harvested at the wound and distal to the site of injury. Harvested tails were fixed in 10% neutral buffered formalin (3800600; Leica Biosystems, Wetzlar, Germany) were cryo-sectioned into 10 µm sections.

### Tissue dissociation for flow cytometry and FIND

Isolated LNs and LVs were washed in 1 mL PBS (21-030-CV; VWR International) on ice. LNs were added in 500 mL of Collagenase D solution (1 mg/mL in PBS; 11088866001; Sigma Aldrich)/ DNase I (40 mg/mL in PBS; 10104159001; Sigma Aldrich)) and incubated for 1 hr at 37°C on rocker/vortex at 300 rpm. LVs were added in 500 mL of Dispase II/Collagenase I mixture (Dispase II (50 mg; 4942078001; Sigma Aldrich), Collagenase I (20 mg; 17-100-017; ThermoFisher Scientific), and BSA (0.1 g; A7906; Sigma Aldrich) in 10 mL DMEM (11039-047; ThermoFisher Scientific). The cell suspension was passed through a 70 µm strainer (07-201-431; ThermoFisher Scientific). Any remaining tissue samples were gently disrupted using the syringe plunger. Filtered cells were centrifuged at 300 g for 5 min at 4°C. Suspended cells were then used for subsequent experiments.

### Flow cytometry

Suspended cells prepared as above were stained for live/dead cell quantification with the Zombie NIR Fixable Viability Kit following with the manufacture’s protocol (1:100 dilution, 423111; BioLegend, San Diego, CA). Subsequently, cells washed with FACs buffer (10 mg/mL BSA (A7906; Sigma Aldrich) in PBS). Antibodies were prepared in the FACs buffer and cells were stained on ice for 30 min in the dark. Information of corresponding laser, concentration, and vendor information of antibodies used for the flow panel in this study was as follows: (i) Live/Dead (BV510, 1:100, 423111; BioLegend), (ii) CD31 (BV605, 1:100, 102416; BioLegend), (iii) PDPN (FITC, 1:100, 156208; BioLegend), (iv) aVHH (APC, 1:100, A01994; GenScript, Piscataway, NJ), (v) CD45 (PE, 1:100, 147712; BioLegend), (vi) CD54 (PE-Cy7, 1:300, 116122; BioLegend), (vii) CD309 (PER-CP-Cy5-5, 1:100, 121918; BioLegend), (viii) CD11b (BV421, 1:100, 101236; BioLegend), (ix) MHCII (BV650, 1:1500, NBP2-00462; Novus Biologicals, Littleton, CO), (x) CD11c (BV786, 1:10, 117335; BioLegend), (xi) CD64 (BV711, 1:100, 139311; BioLegend), and (xii) F4-80 (APC-Cy7, 1:100, 157315; BioLegend). Compensation controls for antibodies were made using UltraComp eBeads™ Compensation Beads (01-2222-42; ThermoFisher Scientific). Data was acquired on the BD FACS Aria III Cell Sorter (BD Biosciences) and analyzed with FlowJo Software. The gating strategy is described in Fig. S8 and Fig. S9. Relevant cell counts are presented in Table S1, Table S2, and Table S3.

### In vivo EdU labeling

During lymphatic injury studies, EdU labeling and its detection was followed with the protocol of Click-iT EdU Cell Proliferation Kit for Imaging Alexa Fluor™ 594 dye (C10639; ThermoFisher Scientific). Briefly, 10 mM of EdU was prepared by diluting EdU with 2mL of DMSO. 25 mg/kg of EdU was injected intradermally into the tail 16 hrs prior to tail harvest.

### EdU detection and Immunofluorescence staining

For antigen retrieval, tail sections were incubated in sodium citrate at 90°C for 30 min. Sections were permeabilized with 0.5% Triton X-100 (X100-5ML; Sigma-Aldrich) in PBS at room temperature for 20 min. EdU were detected with Click-iT Reaction cocktail for 30 min at room temperature protected from the light. Afterwards, DNA was stained with Hoechst solution for 30 min at room temperature in the dark.

After washing the Hoechst solution on tissues with PBS, non-specific binding was blocked with 10% goat serum (G9023; Sigma-Aldrich) in PBS for 1 hour at room temperature. Slides were probed with hamster IgG monoclonal anti-PDPN (1 mg/mL in PBS; ab11936; Abcam, Cambridge, UK) at 1:100 dilution in PBS solution overnight at 4 °C. The following day, the slides were washed with PBS, and incubated in the dark with Alexa Fluor 488-conjugated goat anti-hamster IgG (2 mg/ mL; A21110; ThermoFisher) at 1:200 dilution in PBS solution for 4 hours at room temperature. For nuclei staining and mounting sections, Invitrogen ProLong Gold AntiFade with DAPI (ThermoFisher) was used.

A Zeiss Axio Observer fluorescent microscope was used to image slides after staining, and analysis was performed on high-powered sections (20X objective) with at least 5 high-powered fields (hpf) per location (i.e., wound, or distal) per mouse.

### Single LV ligation lymphatic injury model

To induce lymphatic injury on the mice tail, single LV ligation surgery was performed (*54,79*). Briefly, animals were anesthetized with 5% isoflurane maintained on 2-2.5% of isoflurane during the entire surgery. Prior to surgery, injection of NIR dye was injected intradermally to the tail for the visualization of both the dominant and non-dominant LVs. Animals received incisions 1.6 cm from the base of the tail spanning 80–90% of the circumference of the tail. Non-dominant vessel was left untouched to serve as an internal control. Animals in which LVs were improperly ligated or in which blood vessels were ligated were excluded from the study. For tissue collection, animals were euthanized using CO_2_.

### NIR imaging for monitoring lymphatic function in vivo

NIR lymphatic imaging was performed according to previously published methods (*67*). Before lymphatic vessel imaging, LI-COR IRDye 800 CW (929-70021; LI-COR Biosciences, Lincoln, NE) was diluted in a DMSO to a concentration of 10mg/mL. Then, 10 μL of the dye solution was injected intradermally into the tip of the tail.

The lymphatic vessels imaging was recorded with a customized imaging system consisting of a Lambda LS Xenon arc lamp (LB-LS; Sutter Instrument, Novato, CA), an Olympus MVX-ZB10 microscope (Olympus Corporation, Japan), a 769 nm band-pass excitation filter (49 nm full-width half maximum; FWHM), an 832 nm band-pass emission filter (45 nm FWHM), and an 801.5 nm long-pass dichroic mirror. Images were acquired with a Photometrics Evolve Delta 512 EM-CCD (Teledyne Photometrics, Tucson, AZ). The field of view was centered on the mouse’s tail 7 cm downstream towards the base of the tail from the injection site at the tip of tail. Animals were imaged continuously from the time of injection until 20 minutes post-injection with a 50 ms exposure time and a frame rate of 10 fps. Baseline NIR metrics and tail images were collected in all groups prior to surgery (day 0). For dosage optimization study, NIR functional metrics were again measured after surgery on day 7 prior to euthanasia and tissue collection. When we measured the therapeutic effect of VEGFC mRNA-LNP, NIR functional metrics were measured after surgery on days 7 and 14. Tail volume measurements were taken after surgery on days 3, 7, and 14. Animals were euthanized, and tissue was collected on day 14.

### NIR analysis for quantifying lymphatic function

Analysis of NIR functional metrics was performed during the steady-state period ranging from 5-20 minutes after injection, as defined previously (*74*). Packets of fluorescence were detected by identifying peaks and troughs in the fluorescence signal over time. These measurements were used to calculate previously reported metrics for this model such as packet frequency, amplitude, integral, and transport (*61,74*). All data were normalized to baseline NIR intensity. Sample size for each experiment is included in the corresponding figure caption.

### Quantification of microscopic images

Images of mouse tails were segmented in ImageJ, and the corresponding diameters and lengths were measured. Total tail volume was calculated by truncated cone volume equation for each segment, summed. Absolute tail volume change was calculated by subtracting the corresponding tail measurement obtained in all groups prior to surgery (day 0) and normalized tail volume by dividing by this measurement. Sample size for each experiment is included in the corresponding figure caption.

Subsequently, total LV area, LV perimeter, and LV number per square mm were measured in ImageJ for at least 5 hpf per location (i.e., wound, distal) per mouse. LVs were identified by positive staining for PDPN. LVs were manually selected, and area and perimeter of each selection was measured. The number of LVs was determined as the number of distinct selections per hpf. For EdU staining, lymphatic-specific proliferation was identified by quantifying the colocalization of PDPN and EdU. Specifically, the ImageJ plugin “*JACoP”* (*80*) and the Pearson’s coefficient (*81*) were used. A minimum of five specimens was analyzed per condition/tissue sample. Actual number of samples is included in the corresponding figure caption.

### Statistical Analysis

To compare lymphatic uptake among the lead LEC-specific LNPs, one-way ANOVA was used combined with Robust regression and Outlier removal (ROUT). To compare uptake by LNs and LVs among saline, free aVHH, MC3, and LNP7, two-way ANOVA was used with Tukey’s method to correct for multiple comparisons. To compare uptake among different cell types, ordinary one-way ANOVA was used. To compare the effect of different dosages of VEGFC mRNA in NIR metrics, one-way ANOVA was used with Tukey’s multiple comparisons correction. To compare the effect of different LNPs-loaded with VEGFC mRNA in NIR metrics, mixed-effects analysis was used with Tukey’s multiple comparisons correction. To compare absolute tail volume change, two-way ANOVA with Geisser-Greenhouse correction was used with Tukey’s multiple comparisons correction. To compare the normalized tail volume, a simple linear regression model was used. All histological measurements were compared between groups by nested one-way ANOVA with Tukey’s multiple comparisons test after ROUT to remove outliers within an individual specimen and tissue location. An unpaired *t*-test was used to compare different types of administration. Each data point corresponds to either an independent experiment or the average of each corresponding condition as stated in the figure caption. Data were analyzed using GraphPad Prism 7 (GraphPad Software, San Diego, CA). Reported *p* values are multiplicity adjusted to account for multiple comparisons. For all cases, significance was defined as *p* < 0.05 (*) or *p* < 0.01 (**), *p* < 0.001 (***), or *p* < 0.0001 (****).

## List of Supplementary Materials

### Materials and Methods

Figs. S1 to S10

Tables S1 to S4

Reference (*82*)

## Acknowledgments

We thank Dr. Philip J. Santangelo (Georgia Institute of Technology and Emory University) for providing custom-made aVHH mRNA.

## Funding

American Heart Association Postdoctoral Fellowship grant 829544 (EM) and National Institutes of Health Grant R01HL133216.

## Author contributions

Conceptualization: EM, JED, JBD

Investigation: EM, RC, KJ, ZQ, LNL, YGV, ESE, KP

Funding acquisition: EM, JED, JBD

Project administration: EM, JBD

Supervision: EM, JED, JBD

Writing – original draft: EM

Writing – review & editing: KJ, LNL, ESE, BJT, NP, JED, JBD

## Competing interests

Dahlman, Dixon, Michalaki, and Tamburini are authors on a patent application associated with this work.

## Data and materials availability

All data are available in the main text or the supplementary materials.

## Notes

### Competing Interest Statement

Dixon, Michalaki, Tamburini, and Dahlman are authors on a patent application associated with this work.

