## Supplemental Information for "Lymphatic endothelial cell-targeting lipid nanoparticles delivering VEGFC mRNA improve lymphatic function after injury"

### SUPPLEMENTARY MATERIALS

#### DETAILED MATERIALS AND METHODS

##### LNP formulation

Nucleic acids (mRNA and DNA barcodes) were diluted in 10 mM citrate buffer pH 3, while lipomer, PEG, cholesterol, and helper lipids were diluted in ethanol. The compounds and their molar ratios for the lipid phase of the lead LEC-specific LNPs and MC3-based LNP (MC3) were listed on the Table S4 (*46*). LNPs were formulated by injecting the citrate and lipid phase into a microfluidic device as previously described (Fig. S10) (*46-48*). The flow rates of citrate and lipid phases are 600 μL/min and 200 μL/min respectively. The syringes (Hamilton Company) for injection to the microfluidic device were controlled by syringe pumps (Harvard Apparatus) programmed using FLOWCONTROL™ software (Harvard Apparatus). The weight ratio of lipomer:mRNA in this study was maintained at 10:1.

The ionizable lipids used in this study were stereo-pure lipomers (C12S, C12R, C13S, and C13R were provided from the Dahlman lab), MC3 (555308; MedKoo Biosciences, Inc, Morrisville, NC) and cKK-E12 (BP-29590; BroadPharm, San Diego, CA). All PEG, cholesterol, and helper lipids were purchased from Avanti Polar Lipids (Alabaster, AL).

##### LNP characterization

After LNPs were diluted in sterile 1X PBS to a concentration of ~0.06 μg/mL, the hydrodynamic diameter (nm), polydispersity (PD), and polydispersity index (PDI) of LNP were measured using dynamic light scattering (DLS) using DynaPro Plate Reader II (Wyatt Technology). LNPs were included in the experimental design if they met all of the following criteria: (i) diameter >20 nm, (ii) diameter <150 nm, (iii) correlation function with one inflection point, and (iv) PDI <0.4.

Quality-controlled LNPs were dialyzed with 1X PBS using dialysis cassettes with membrane (87735 and 87734; ThermoFisher Scientific, Waltham, CA) for 90 minutes. Dialyzed LNPs were sterile-filtered with a 0.22-µm filter (371-2115-OEM; Foxx Life Sciences, Londonderry, NH).

The nucleic acid concentration of the filtered LNPs was measured using Nanodrop (ND-ONE-W; ThermoFisher Scientific). To quantify the nucleic acid concentration inside the LNP, the LNP was loaded in a 96-well plate (675097; Greiner Bio-One), and the RiboGreen assay was performed following manufacturer's protocol (R11490; Invitrogen, Waltham, MA).

**SANDS (Species agnostic nanoparticle delivery screening)**

In the screening of LNPs to identify LEC-specific LNPs, SANDS was conducted as previously described (*48*). Briefly, 150 LNPs with varying lipid compositions were formulated with 56-nucleotide-long ssDNA sequences serving as DNA barcodes and aVHH mRNA. Each ssDNA sequence contained a unique 8-bp barcode sequence in its center, and these sequences were purchased from Integrated DNA Technologies. Quality-controlled LNPs were screened in female C57Bl/6 mice via intradermal injection in each paw of mice at a dosage of 1.5 mg/kg, and aVHH+ lymph nodes (ALN, BLN, and PLN) were isolated and sequenced using Illumina Miniseq with primers from Nextera XT adapter sequences. Lymphatic uptake was quantified based on the normalized barcode counts for each LN using a custom Python-based tool.

**Cell culturing and LNP transfection**

Human dermal LECs were isolated from human foreskin tissue following the protocol published by Rogic and coworkers (*82*). The cells were seeded in a 24-well plate (353047; ThermoFisher Scientific) at a density of 15,000 cells/ well. After 24 hrs, LNP7 was added with a total aVHH mRNA dose of 4, 20, or 100 ng in eight separate wells (*46*). 6 hrs post-transfection, media was removed and replaced with fresh media. Cells cultured with physiological endothelial basal medium (EBM; CC-3121; Lonza, Switzerland) with recommended supplements and 10% DMSO (D2650; Sigma Aldrich) served as negative and positive controls, respectively.

**In Vitro toxicity study of LNP7 using Live/Dead staining and alamarBlue assay**

The viability of monolayers after treatment was determined using a viability kit (L3224; Thermo Fisher) to distinguish live (Calcein-AM) and dead cells (Ethidium homodimer-1). Staining was performed following the manufacturer's instructions; cells were incubated with calcein-AM and ethidium homodimer-1 for 20 minutes at 37°C. Then, monolayers were rinsed with PBS before imaging on an inverted microscope (AxioObserver.Z1; Zeiss). Tile images of individual wells were acquired using a 1x tube lens and a 2.5x objective (Plan-Neofluar 2.5x/0.075Pol). Zen Black software was used to stitch tile images with a 10% overlap to reconstruct the image of the entire well. Images were then processed using ImageJ (NIH), where the "watershed" function was utilized to segment individual cells. Following processing, the "analyze particle" function in ImageJ was used to count individual cells in the live (green) and dead (red) channels, with thresholds set to 0-25 and 0-80, respectively. Viability was further confirmed using alamarBlue cell viability reagent (DAL1025; ThermoFisher). According to the manufacturer's instructions, 1X alamarBlue reagent was added to the cell media of monolayers after treatment and incubated for 6 hours at 37°C. Fluorescence intensity was measured with an excitation of 530nm and emission of 590nm using a plate reader (Synergy H4; BioTek). The average background fluorescence was subtracted, and individual fluorescence intensity per well was reported.

**FIGURES**

Screening 1

| Name | Lipomer | Cholesterol | PEG | Helper Lipid | Lipomer | Cholesterol | PEG | Helper Lipid |
|---|---|---|---|---|---|---|---|---|
| LNP 1 | cKK-E12 | Cholesterol | C14PEG2K | DSPC | 30 | 30 | 2.5 | 37.5 |
| LNP 2 | cKK-E12 | Cholesterol | C14PEG2K | DSPC | 35 | 18 | 2.5 | 44.5 |
| LNP 3 | cKK-E12 | Cholesterol | C14PEG2K | DSPC | 45 | 42 | 2.5 | 10.5 |
| LNP 4 | cKK-E12 | Cholesterol | C14PEG2K | DSPC | 50 | 35 | 2.5 | 12.5 |
| LNP 5 | cKK-E12 | Cholesterol | C14PEG2K | DSPC | 52.5 | 15 | 2.5 | 30 |
| LNP 6 | cKK-E12 | Cholesterol | C14PEG2K | DSPC | 57.5 | 20 | 2.5 | 20 |
| LNP 7 | cKK-E12 | Cholesterol | C14PEG2K | DOPE | 30 | 30 | 2.5 | 37.5 |
| LNP 8 | cKK-E12 | Cholesterol | C14PEG2K | DOPE | 35 | 18 | 2.5 | 44.5 |
| LNP 9 | cKK-E12 | Cholesterol | C14PEG2K | DOPE | 45 | 42 | 2.5 | 10.5 |
| LNP 10 | cKK-E12 | Cholesterol | C14PEG2K | DOPE | 50 | 35 | 2.5 | 12.5 |
| LNP 11 | cKK-E12 | Cholesterol | C14PEG2K | DOPE | 52.5 | 15 | 2.5 | 30 |
| LNP 12 | cKK-E12 | Cholesterol | C14PEG2K | DOPE | 57.5 | 20 | 2.5 | 20 |
| LNP 13 | cKK-E12 | Cholesterol | C14PEG2K | 18:1 CAP PE | 30 | 30 | 2.5 | 37.5 |
| LNP 14 | cKK-E12 | Cholesterol | C14PEG2K | 18:1 CAP PE | 35 | 18 | 2.5 | 44.5 |
| LNP 15 | cKK-E12 | Cholesterol | C14PEG2K | 18:1 CAP PE | 45 | 42 | 2.5 | 10.5 |
| LNP 16 | cKK-E12 | Cholesterol | C14PEG2K | 18:1 CAP PE | 50 | 35 | 2.5 | 12.5 |
| LNP 17 | cKK-E12 | Cholesterol | C14PEG2K | 18:1 CAP PE | 52.5 | 15 | 2.5 | 30 |
| LNP 18 | cKK-E12 | Cholesterol | C14PEG2K | 18:1 CAP PE | 57.5 | 20 | 2.5 | 20 |
| LNP 19 | cKK-E12 | Cholesterol | C14PEG2K | 18:0 DDAB | 30 | 30 | 2.5 | 37.5 |
| LNP 20 | cKK-E12 | Cholesterol | C14PEG2K | 18:0 DDAB | 35 | 18 | 2.5 | 44.5 |
| LNP 21 | cKK-E12 | Cholesterol | C14PEG2K | 18:0 DDAB | 45 | 42 | 2.5 | 10.5 |
| LNP 22 | cKK-E12 | Cholesterol | C14PEG2K | 18:0 DDAB | 50 | 35 | 2.5 | 12.5 |
| LNP 23 | cKK-E12 | Cholesterol | C14PEG2K | 18:0 DDAB | 52.5 | 15 | 2.5 | 30 |
| LNP 24 | cKK-E12 | Cholesterol | C14PEG2K | 18:0 DDAB | 57.5 | 20 | 2.5 | 20 |
| LNP 25 | cKK-E12 | Cholesterol | C14PEG2K | DOTAP | 30 | 30 | 2.5 | 37.5 |
| LNP 26 | cKK-E12 | Cholesterol | C14PEG2K | DOTAP | 35 | 18 | 2.5 | 44.5 |
| LNP 27 | cKK-E12 | Cholesterol | C14PEG2K | DOTAP | 45 | 42 | 2.5 | 10.5 |
| LNP 28 | cKK-E12 | Cholesterol | C14PEG2K | DOTAP | 50 | 35 | 2.5 | 12.5 |
| LNP 29 | cKK-E12 | Cholesterol | C14PEG2K | DOTAP | 52.5 | 15 | 2.5 | 30 |
| LNP 30 | cKK-E12 | Cholesterol | C14PEG2K | DOTAP | 57.5 | 20 | 2.5 | 20 |

Screening 2

| Name | Lipomer | Cholesterol | PEG | Helper Lipid | Lipomer | Cholesterol | PEG | Helper Lipid |
|---|---|---|---|---|---|---|---|---|
| LNP 31 | KB11 | Cholesterol | C14PEG | DOTAP | 45.0 | 38.8 | 2.5 | 13.7 |
| LNP 32 | KB11 | Cholesterol | C14PEG | DOTAP | 50.0 | 35.0 | 2.5 | 12.5 |
| LNP 33 | KB11 | 20a-OH | C14PEG | DOTAP | 40.0 | 42.7 | 2.5 | 14.8 |
| LNP 34 | KB11 | 20a-OH | C14PEG | DOTAP | 45.0 | 38.8 | 2.5 | 13.7 |
| LNP 35 | KB11 | 20a-OH | C14PEG | DOTAP | 45.0 | 41.5 | 1.5 | 12.0 |
| LNP 36 | KB12 | Cholesterol | C14PEG | DOTAP | 35.0 | 46.5 | 2.5 | 16.0 |
| LNP 37 | KB12 | Cholesterol | C14PEG | DOTAP | 35.0 | 43.5 | 1.5 | 20.0 |
| LNP 38 | | | | | | | | |
| LNP 39 | | | | | | | | |
| LNP 40 | KB11 | Cholesterol | C14PEG | DOPE | 45.0 | 38.8 | 2.5 | 13.7 |
| LNP 41 | KB11 | Cholesterol | C14PEG | DOPE | 50.0 | 35.0 | 2.5 | 12.5 |
| LNP 42 | KB11 | 20a-OH | C14PEG | DOPE | 40.0 | 42.7 | 2.5 | 14.8 |
| LNP 43 | KB11 | 20a-OH | C14PEG | DOPE | 45.0 | 38.8 | 2.5 | 13.7 |
| LNP 44 | KB11 | 20a-OH | C14PEG | DOPE | 45.0 | 41.5 | 1.5 | 12.0 |
| LNP 45 | KB12 | Cholesterol | C14PEG | DOPE | 35.0 | 46.5 | 2.5 | 16.0 |
| LNP 46 | KB12 | Cholesterol | C14PEG | DOPE | 35.0 | 43.5 | 1.5 | 20.0 |
| LNP 47 | KB12 | 20a-OH | C14PEG | DOPE | 35.0 | 46.5 | 2.5 | 16.0 |
| LNP 48 | KB12 | 20a-OH | C14PEG | DOPE | 45.0 | 38.8 | 2.5 | 13.7 |
| LNP 49 | KB12 | 20a-OH | C14PEG | DOPE | 50.0 | 35.0 | 2.5 | 12.5 |
| LNP 50 | KB12 | 20a-OH | C14PEG | DOPE | 35.0 | 43.5 | 1.5 | 20.0 |
| LNP 51 | KB12 | 20a-OH | C14PEG | DOPE | 40.0 | 42.5 | 1.5 | 16.0 |
| LNP 52 | KB12 | 20a-OH | C14PEG | DOTAP | 35.0 | 46.5 | 2.5 | 16.0 |
| LNP 53 | KB12 | 20a-OH | C14PEG | DOTAP | 45.0 | 38.8 | 2.5 | 13.7 |
| LNP 54 | KB12 | 20a-OH | C14PEG | DOTAP | 50.0 | 35.0 | 2.5 | 12.5 |
| LNP 55 | KB12 | 20a-OH | C14PEG | DOTAP | 35.0 | 43.5 | 1.5 | 20.0 |
| LNP 56 | KB12 | 20a-OH | C14PEG | DOTAP | 40.0 | 42.5 | 1.5 | 16.0 |
| LNP 57 | KB15 | Cholesterol | C14PEG | DOPE | 35.0 | 46.5 | 2.5 | 16.0 |
| LNP 58 | KB15 | Cholesterol | C14PEG | DOPE | 40.0 | 42.7 | 2.5 | 14.8 |
| LNP 59 | KB15 | Cholesterol | C14PEG | DOPE | 45.0 | 38.8 | 2.5 | 13.7 |
| LNP 60 | KB15 | Cholesterol | C14PEG | DOPE | 50.0 | 35.0 | 2.5 | 12.5 |
| LNP 61 | KB15 | Cholesterol | C14PEG | DOPE | 40.0 | 42.5 | 1.5 | 16.0 |
| LNP 62 | KB15 | 20a-OH | C14PEG | DOPE | 35.0 | 46.5 | 2.5 | 16.0 |
| LNP 63 | KB15 | 20a-OH | C14PEG | DOPE | 40.0 | 42.7 | 2.5 | 14.8 |
| LNP 64 | KB15 | 20a-OH | C14PEG | DOPE | 45.0 | 38.8 | 2.5 | 13.7 |
| LNP 65 | KB15 | 20a-OH | C14PEG | DOPE | 35.0 | 43.5 | 1.5 | 20.0 |
| LNP 66 | KB15 | Cholesterol | C14PEG | DOTAP | 35.0 | 46.5 | 2.5 | 16.0 |
| LNP 67 | KB15 | Cholesterol | C14PEG | DOTAP | 40.0 | 42.7 | 2.5 | 14.8 |
| LNP 68 | KB15 | Cholesterol | C14PEG | DOTAP | 45.0 | 38.8 | 2.5 | 13.7 |
| LNP 69 | KB15 | Cholesterol | C14PEG | DOTAP | 50.0 | 35.0 | 2.5 | 12.5 |
| LNP 70 | KB15 | Cholesterol | C14PEG | DOTAP | 40.0 | 42.5 | 1.5 | 16.0 |
| LNP 71 | KB15 | 20a-OH | C14PEG | DOTAP | 35.0 | 46.5 | 2.5 | 16.0 |
| LNP 72 | KB15 | 20a-OH | C14PEG | DOTAP | 40.0 | 42.7 | 2.5 | 14.8 |
| LNP 73 | KB15 | 20a-OH | C14PEG | DOTAP | 45.0 | 38.8 | 2.5 | 13.7 |
| LNP 74 | KB16 | Cholesterol | C14PEG | DOPE | 40.0 | 42.7 | 2.5 | 14.8 |
| LNP 75 | KB16 | Cholesterol | C14PEG | DOPE | 40.0 | 42.5 | 1.5 | 16.0 |
| LNP 76 | KB16 | 20a-OH | C14PEG | DOPE | 35.0 | 46.5 | 2.5 | 16.0 |
| LNP 77 | KB16 | 20a-OH | C14PEG | DOPE | 40.0 | 42.7 | 2.5 | 14.8 |
| LNP 78 | KB16 | 20a-OH | C14PEG | DOPE | 45.0 | 38.8 | 2.5 | 13.7 |
| LNP 79 | KB16 | 20a-OH | C14PEG | DOPE | 50.0 | 35.0 | 2.5 | 12.5 |
| LNP 80 | KB16 | 20a-OH | C14PEG | DOPE | 45.0 | 41.5 | 1.5 | 12.0 |
| LNP 81 | KB16 | Cholesterol | C14PEG | DOTAP | 40.0 | 42.7 | 2.5 | 14.8 |
| LNP 82 | KB16 | Cholesterol | C14PEG | DOTAP | 40.0 | 42.5 | 1.5 | 16.0 |
| LNP 83 | KB15 | 20a-OH | C14PEG | DOTAP | 35.0 | 43.5 | 1.5 | 20.0 |
| LNP 84 | KB16 | 20a-OH | C14PEG | DOTAP | 35.0 | 46.5 | 2.5 | 16.0 |
| LNP 85 | KB16 | 20a-OH | C14PEG | DOTAP | 40.0 | 42.7 | 2.5 | 14.8 |
| LNP 86 | KB16 | 20a-OH | C14PEG | DOTAP | 45.0 | 38.8 | 2.5 | 13.7 |
| LNP 87 | KB16 | 20a-OH | C14PEG | DOTAP | 50.0 | 35.0 | 2.5 | 12.5 |
| LNP 88 | KB16 | 20a-OH | C14PEG | DOTAP | 45.0 | 41.5 | 1.5 | 12.0 |
| TAACTCGG | | | | | | | | |
| TGCCTTGA | | | | | | | | |
| CCAGAGTA | | | | | | | | |

Screening 3

| Name | Lipomer | Cholesterol | PEG | Helper Lipid | Lipomer | Cholesterol | PEG | Helper Lipid |
|---|---|---|---|---|---|---|---|---|
| LNP 91 | cKK-E12 | Cholesterol | C14PEG2K | DOPE | 45 | 44 | 2 | 9 |
| LNP 92 | cKK-E12 | Cholesterol | C14PEG2K | DOPE | 35 | 46.5 | 2.5 | 16 |
| LNP 93 | cKK-E12 | Cholesterol | C14PEG2K | DOPE | 50 | 35 | 2.5 | 12.5 |
| LNP 94 | cKK-E12 | Cholesterol | C14PEG2K | DOPE | 30 | 30 | 1 | 39 |
| LNP 95 | cKK-E12 | Cholesterol | C14PEG2K | DOPE | 35 | 18 | 2.5 | 44.5 |
| LNP 96 | cKK-E12 | Cholesterol | C14PEG2K | DOTAP | 45 | 44 | 2 | 9 |
| LNP 97 | cKK-E12 | Cholesterol | C14PEG2K | DOTAP | 35 | 46.5 | 2.5 | 16 |
| LNP 98 | cKK-E12 | Cholesterol | C14PEG2K | DOTAP | 50 | 35 | 2.5 | 12.5 |
| LNP 99 | cKK-E12 | Cholesterol | C14PEG2K | DOTAP | 30 | 30 | 1 | 39 |
| LNP 100 | cKK-E12 | Cholesterol | C14PEG2K | DOTAP | 35 | 18 | 2.5 | 44.5 |
| LNP 101 | cKK-E12 | Cholesterol | C14PEG2K | DOTMA | 45 | 44 | 2 | 9 |
| LNP 102 | cKK-E12 | Cholesterol | C14PEG2K | DOTMA | 35 | 46.5 | 2.5 | 16 |
| LNP 103 | cKK-E12 | Cholesterol | C14PEG2K | DOTMA | 50 | 35 | 2.5 | 12.5 |
| LNP 104 | cKK-E12 | Cholesterol | C14PEG2K | DOTMA | 30 | 30 | 1 | 39 |
| LNP 105 | cKK-E12 | Cholesterol | C14PEG2K | DOTMA | 35 | 18 | 2.5 | 44.5 |
| LNP 106 | cKK-E12 | 20a-OH | C14PEG2K | DOPE | 45 | 44 | 2 | 9 |
| LNP 107 | cKK-E12 | 20a-OH | C14PEG2K | DOPE | 35 | 46.5 | 2.5 | 16 |
| LNP 108 | cKK-E12 | 20a-OH | C14PEG2K | DOPE | 50 | 35 | 2.5 | 12.5 |
| LNP 109 | cKK-E12 | 20a-OH | C14PEG2K | DOPE | 30 | 30 | 1 | 39 |
| LNP 110 | cKK-E12 | 20a-OH | C14PEG2K | DOPE | 35 | 18 | 2.5 | 44.5 |
| LNP 111 | cKK-E12 | 20a-OH | C14PEG2K | DOTAP | 45 | 44 | 2 | 9 |
| LNP 112 | cKK-E12 | 20a-OH | C14PEG2K | DOTAP | 35 | 46.5 | 2.5 | 16 |
| LNP 113 | cKK-E12 | 20a-OH | C14PEG2K | DOTAP | 50 | 35 | 2.5 | 12.5 |
| LNP 114 | cKK-E12 | 20a-OH | C14PEG2K | DOTAP | 30 | 30 | 1 | 39 |
| LNP 115 | cKK-E12 | 20a-OH | C14PEG2K | DOTAP | 35 | 18 | 2.5 | 44.5 |
| LNP 116 | cKK-E12 | 20a-OH | C14PEG2K | DOTMA | 45 | 44 | 2 | 9 |
| LNP 117 | cKK-E12 | 20a-OH | C14PEG2K | DOTMA | 35 | 46.5 | 2.5 | 16 |
| LNP 118 | cKK-E12 | 20a-OH | C14PEG2K | DOTMA | 50 | 35 | 2.5 | 12.5 |
| LNP 119 | cKK-E12 | 20a-OH | C14PEG2K | DOTMA | 30 | 30 | 1 | 39 |
| LNP 120 | cKK-E12 | 20a-OH | C14PEG2K | DOTMA | 35 | 18 | 2.5 | 44.5 |
| LNP 121 | cKK-E12 | 20a-OH | C18PEG2K | DOPE | 45 | 44 | 2 | 9 |
| LNP 122 | cKK-E12 | 20a-OH | C18PEG2K | DOPE | 35 | 46.5 | 2.5 | 16 |
| LNP 123 | cKK-E12 | 20a-OH | C18PEG2K | DOPE | 50 | 35 | 2.5 | 12.5 |
| LNP 124 | cKK-E12 | 20a-OH | C18PEG2K | DOPE | 30 | 30 | 1 | 39 |
| LNP 125 | cKK-E12 | 20a-OH | C18PEG2K | DOPE | 35 | 18 | 2.5 | 44.5 |
| LNP 126 | cKK-E12 | 20a-OH | C18PEG2K | DOTAP | 45 | 44 | 2 | 9 |
| LNP 127 | cKK-E12 | 20a-OH | C18PEG2K | DOTAP | 35 | 46.5 | 2.5 | 16 |
| LNP 128 | cKK-E12 | 20a-OH | C18PEG2K | DOTAP | 50 | 35 | 2.5 | 12.5 |
| LNP 129 | cKK-E12 | 20a-OH | C18PEG2K | DOTAP | 30 | 30 | 1 | 39 |
| LNP 130 | cKK-E12 | 20a-OH | C18PEG2K | DOTAP | 35 | 18 | 2.5 | 44.5 |
| LNP 131 | cKK-E12 | 20a-OH | C18PEG2K | DOTMA | 45 | 44 | 2 | 9 |
| LNP 132 | cKK-E12 | 20a-OH | C18PEG2K | DOTMA | 35 | 46.5 | 2.5 | 16 |
| LNP 133 | cKK-E12 | 20a-OH | C18PEG2K | DOTMA | 50 | 35 | 2.5 | 12.5 |
| LNP 134 | cKK-E12 | 20a-OH | C18PEG2K | DOTMA | 30 | 30 | 1 | 39 |
| LNP 135 | cKK-E12 | 20a-OH | C18PEG2K | DOTMA | 35 | 18 | 2.5 | 44.5 |
| LNP 136 | cKK-E12 | Cholesterol | C18PEG2K | DOPE | 45 | 44 | 2 | 9 |
| LNP 137 | cKK-E12 | Cholesterol | C18PEG2K | DOPE | 35 | 46.5 | 2.5 | 16 |
| LNP 138 | cKK-E12 | Cholesterol | C18PEG2K | DOPE | 50 | 35 | 2.5 | 12.5 |
| LNP 139 | cKK-E12 | Cholesterol | C18PEG2K | DOPE | 30 | 30 | 1 | 39 |
| LNP 140 | cKK-E12 | Cholesterol | C18PEG2K | DOPE | 35 | 18 | 2.5 | 44.5 |
| LNP 141 | cKK-E12 | Cholesterol | C18PEG2K | DOTAP | 45 | 44 | 2 | 9 |
| LNP 142 | cKK-E12 | Cholesterol | C18PEG2K | DOTAP | 35 | 46.5 | 2.5 | 16 |
| LNP 143 | cKK-E12 | Cholesterol | C18PEG2K | DOTAP | 50 | 35 | 2.5 | 12.5 |
| LNP 144 | cKK-E12 | Cholesterol | C18PEG2K | DOTAP | 30 | 30 | 1 | 39 |
| LNP 145 | cKK-E12 | Cholesterol | C18PEG2K | DOTAP | 35 | 18 | 2.5 | 44.5 |
| LNP 146 | cKK-E12 | Cholesterol | C18PEG2K | DOTMA | 45 | 44 | 2 | 9 |
| LNP 147 | cKK-E12 | Cholesterol | C18PEG2K | DOTMA | 35 | 46.5 | 2.5 | 16 |
| LNP 148 | cKK-E12 | Cholesterol | C18PEG2K | DOTMA | 50 | 35 | 2.5 | 12.5 |
| LNP 149 | cKK-E12 | Cholesterol | C18PEG2K | DOTMA | 30 | 30 | 1 | 39 |
| LNP 150 | cKK-E12 | Cholesterol | C18PEG2K | DOTMA | 35 | 18 | 2.5 | 44.5 |

**Fig. S1. Detailed outline of total 150 LNPs with varying composition for screenings**

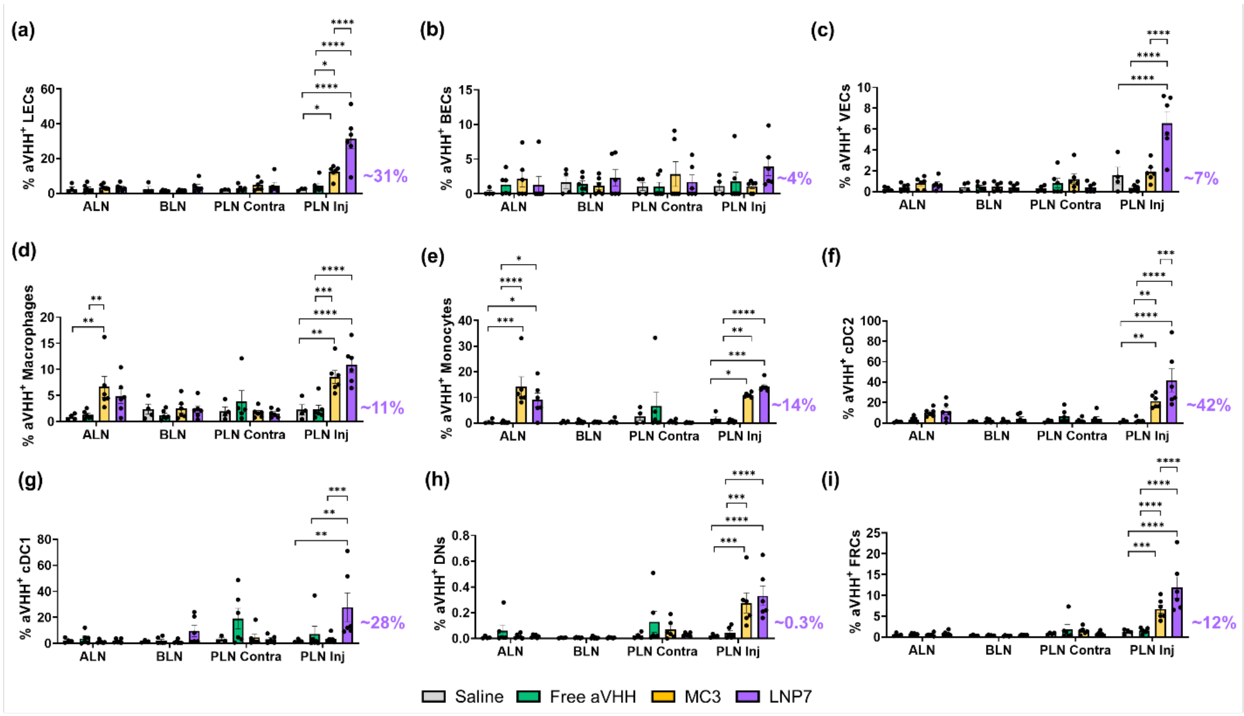


**Fig. S2. Lymphatic-specific uptake of LNP7.** Percentage of aVHH$^+$: **(a)** LECs, **(b)** BECs, **(c)** VECs, **(d)** Macrophages, **(e)** Monocytes, **(f)** cDC2, **(g)** cDC1, **(h)** DNs, and **(i)** FRCs from ALN, BLN, PLN Contra (non-injected), and PLN Inj that have successfully taken up saline (gray), free aVHH (green), MC3 (gold) and LNP7 (purple). Each data point corresponds to an independent experiment (ALN: $N_{Saline} = 4$, $N_{Free\ aVHH} = 6$, $N_{MC3} = 6$, and $N_{LNP7} = 6$; BLN: $N_{Saline} = 4$, $N_{Free\ aVHH} = 6$, $N_{MC3} = 6$, and $N_{LNP7} = 6$, PLN Contra: $N_{Saline} = 4$, $N_{Free\ aVHH} = 6$, $N_{MC3} = 6$, and $N_{LNP7} = 6$; PLN Inj: $N_{Saline} = 4$, $N_{Free\ aVHH} = 6$, $N_{MC3} = 6$, and $N_{LNP7} = 6$), and error bars represent the standard error of the mean. Solid lines above plots indicate a pairwise comparison for significance using a two-way ANOVA with Tukey's multiple comparisons test with $p < 0.05$ (*), $p < 0.01$ (**), $p < 0.001$ (***), and $p < 0.0001$ (****).

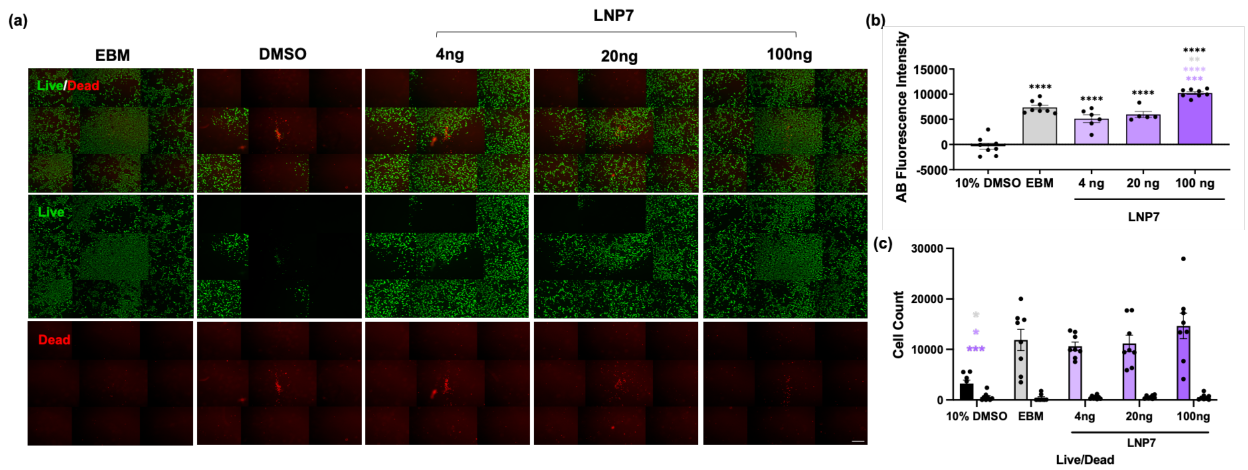


**Fig. S3. LNP7 does not significantly lower the viability of LECs. (a)** Live (Calcein-AM, green) and Dead (TOTO-3, red) staining of human LEC monolayers after treatment with controls or LNP7. (Scale bar = 200 μm.) Contrast was enhanced post-acquisition equally in all image panels for ease of viewing. Quantification of **(b)** Alamar Blue fluorescence intensity (metabolic activity) and **(c)** Live/Dead staining. **(b)** Color-coordinated asterisks indicate comparison with the corresponding treatment using one-way ANOVA with Tukey's multiple comparisons test and robust regression and outlier removal (ROUT) method to identify and remove outliers with $p < 0.01$ (**), $p < 0.001$ (***), and $p < 0.0001$ (****). **(c)** Cell count of Live (left bar) and Dead (right bar) positive stain. Asterisks indicate significant differences by Sídák's multiple comparisons test with $p < 0.05$ (*), $p < 0.01$ (**), $p < 0.001$ (***), and $p < 0.0001$ (****).

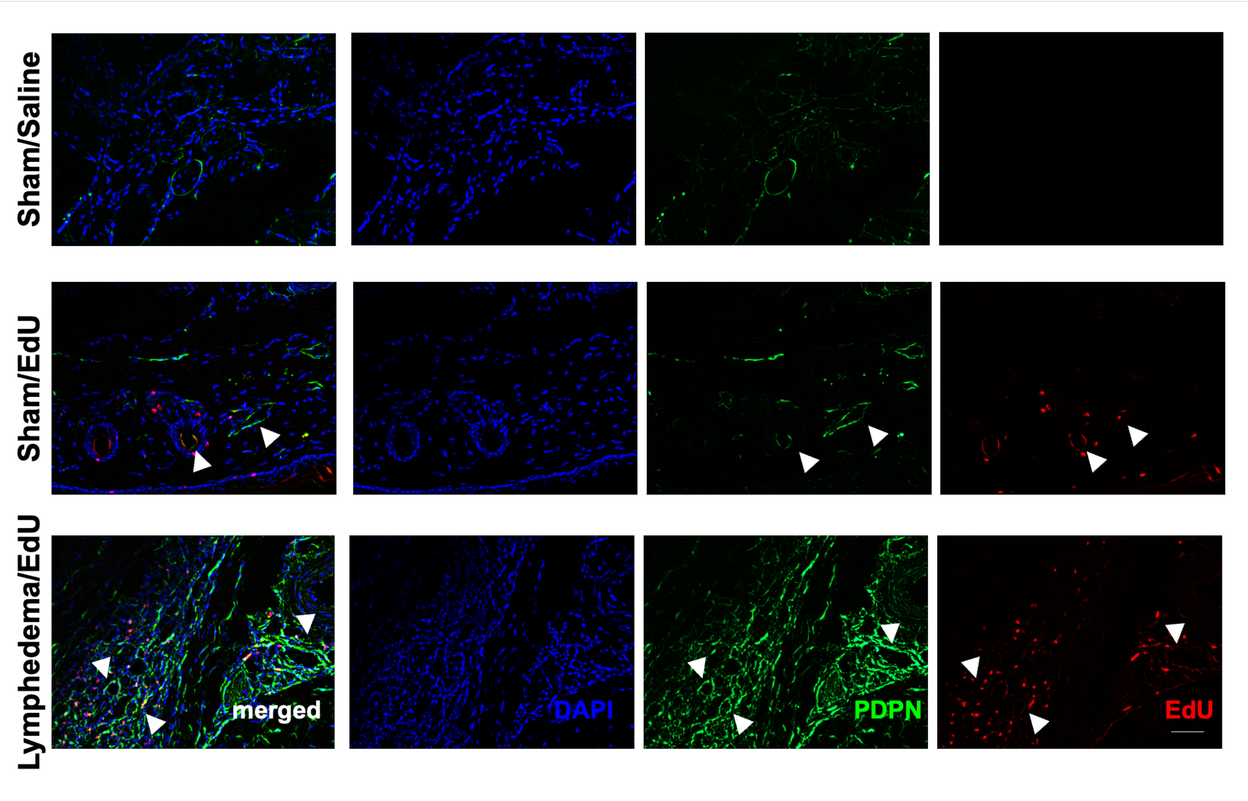


**Fig. S4. IHC controls in tail sections.** Immunofluorescence micrographs of tail LV segments for sham mice injected with saline (negative control), sham mice injected with EdU, and lymphedema mice injected with EdU 7 days post-surgery for merged, DAPI (blue), PDPN (green), and EdU (red). (20x objective; Scale bar = 50 μm.) Arrows indicate EdU and PDPN double positive LEC. The contrast was enhanced post-acquisition equally in all image panels for ease of viewing.

| | Characterization | | |
|---|---|---|---|
| | **mRNA Cargo** | **aVHH** | **VEGFC** |
| MC3 | Diameter (nm) | 82 | 85 |
| | PDI | 0.10 | 0.29 |
| | Encapsulation Efficiency (%) | 82 | 67 |
| | Total mRNA Concentration (μg/mL) | 62 | 41 |
| | Encapsulated mRNA Concentration (μg/mL) | 51 | 28 |
| LNP7 | Diameter (nm) | 81 | 81 |
| | PDI | 0.17 | 0.15 |
| | Encapsulation Efficiency (%) | 65 | 43 |
| | Total mRNA Concentration (μg/mL) | 50 | 38 |
| | Encapsulated mRNA Concentration (μg/mL) | 33 | 17 |

**Fig. S5. Characterization of MC3 and LNP7 loaded with different mRNA cargos, namely aVHH and VEGFC.** Hydrodynamic diameter (nm), PDI, encapsulation efficiency (%), total mRNA concentration (mg/mL), encapsulated mRNA concentration (mg/mL) of MC3 and LNP7 loaded with aVHH mRNA and VEGFC mRNA.

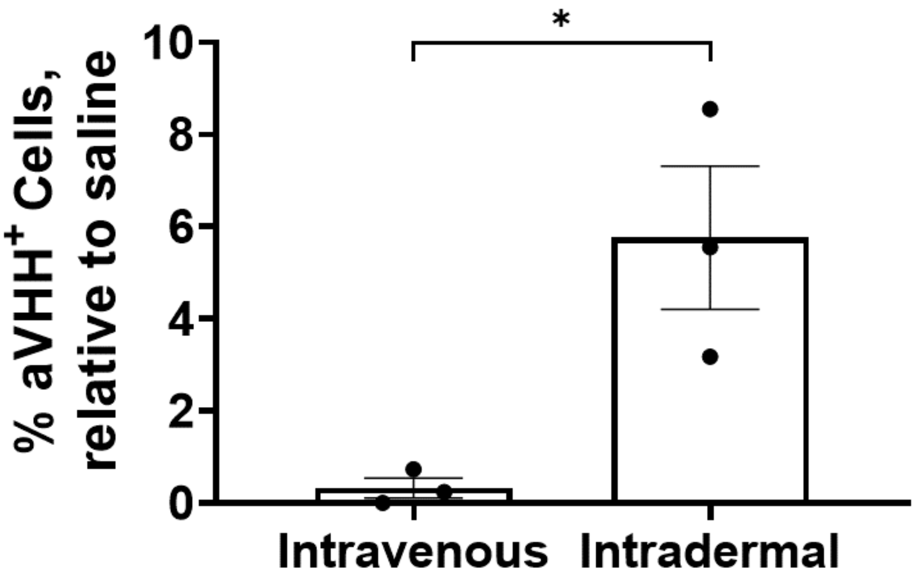


**Fig. S6. Effect of the route of administration on LNP delivery.** Percentage of aVHH$^+$ cells (after gating for Live/Dead, CD31$^+$/PDPN$^+$, and relative to saline (meaning, the saline measurement was subtracted from the corresponding uptake)) in PLN that have successfully taken up the LNP cargo after intravenous and intradermal injections during the LNP screening study. Each data point corresponds to an independent experiment ($N_{Intravenous}$ = 3 and $N_{Intradermal}$ = 3), and error bars represent the standard error of the mean. The solid line above plots indicates a pairwise comparison for significance using an unpaired *t*-test with $p < 0.05$ (*).

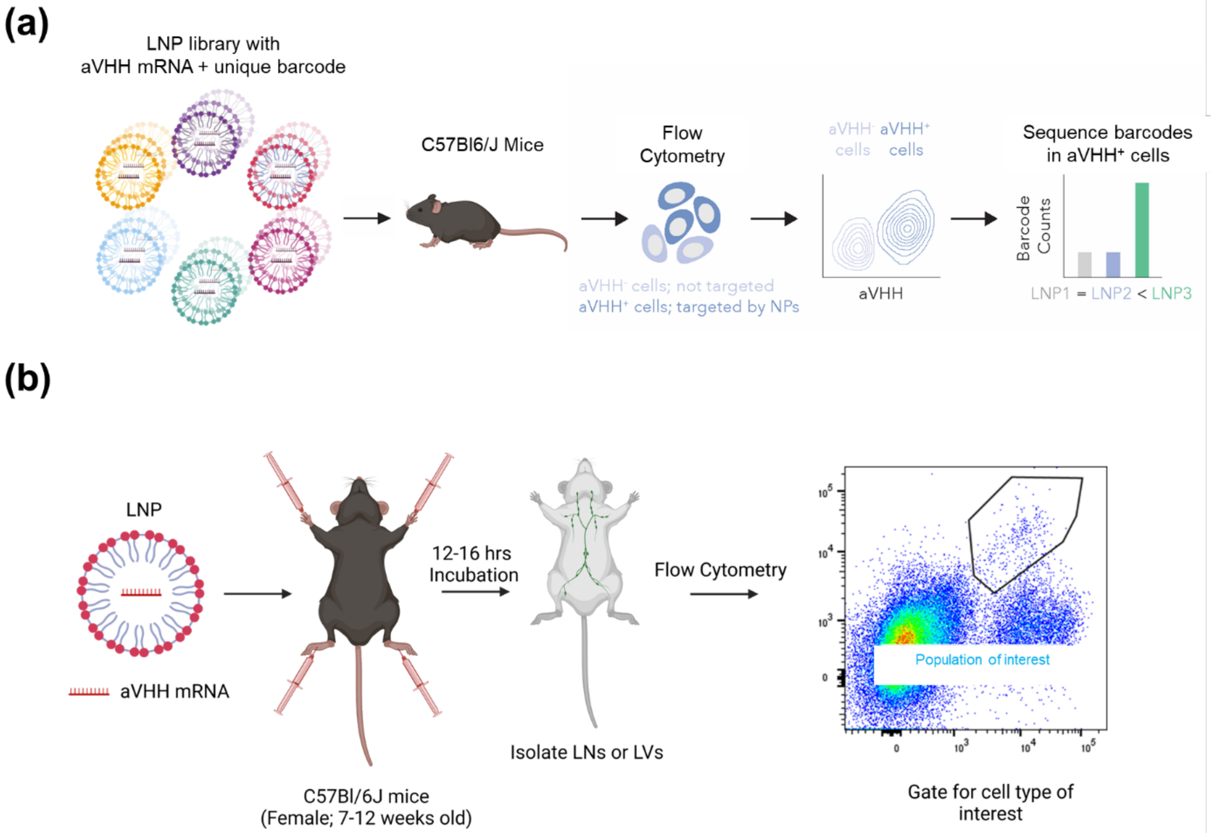


**Fig. S7. Engineer LEC-targeting LNPs. (a)** LNP libraries with aVHH mRNA and unique DNA barcode were administered to C57Bl6/J mice. aVHH$^{+}$ cells were isolated by FACS and DNA barcodes were sequenced. **(b)** LNPs were intradermally injected in C57Bl/6J mice and 12-16 hrs later the downstream LNs and LVs were collected for flow cytometry. Control animals are ID injected with appropriate volumes of saline.

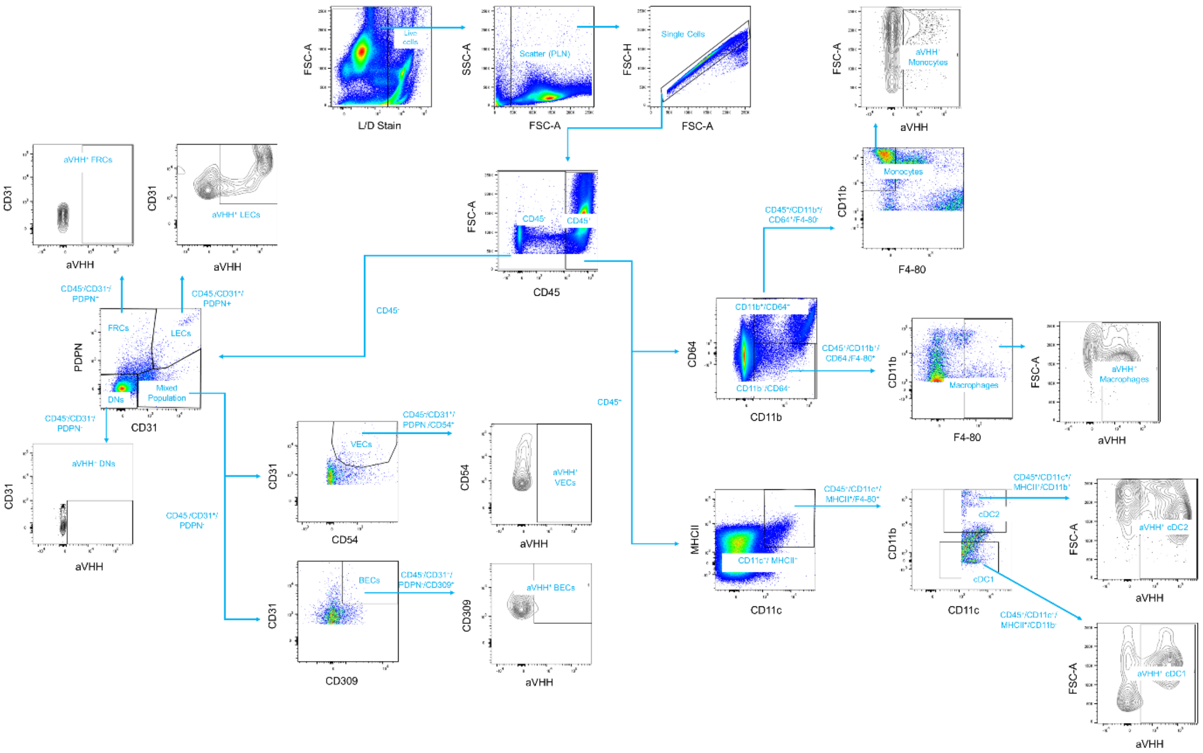

**Fig. S8. Representative gating strategies for FACS for various cell populations present in LNs.** Identifying the following cell populations via appropriate gating as seen above (after Live/Dead gating); LECs: CD45$^-$/CD31$^+$/PDPN$^+$, VECs: CD45$^-$/CD31$^+$/PDPN$^-$/CD54$^+$, BECs: CD45$^-$/CD31$^+$/PDPN$^-$/CD309$^+$, FRCs: CD45$^-$/CD31$^-$/PDPN$^+$, DNs: CD45$^-$/CD31$^-$/PDPN$^-$, Monocytes: CD45$^+$/CD11b$^+$/CD64$^+$/F4-80$^-$, Macrophages: CD45$^+$/CD11b$^+$/CD64$^-$/F4-80$^+$, cDC2: CD45$^+$/CD11c$^+$/MHCII$^+$/CD11b$^+$, and cDC1: CD45$^+$/CD11c$^+$/MHCII$^+$/CD11b$^-$.

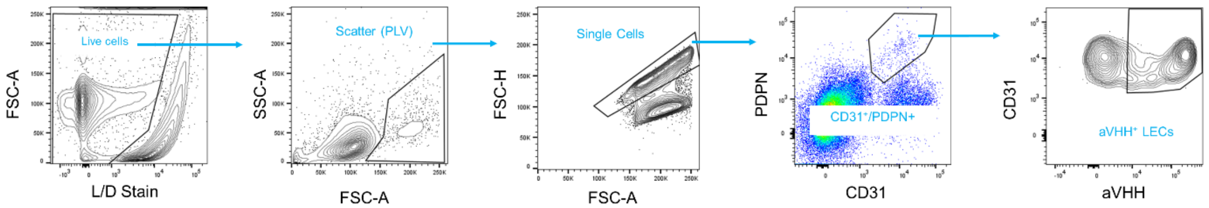


**Fig. S9. Representative gating strategies for FACS for LECs in LVs.** Identifying the LEC population via appropriate gating (after Live/Dead gating) as seen above; LECs: $CD31^{+}/PDPN^{+}$.

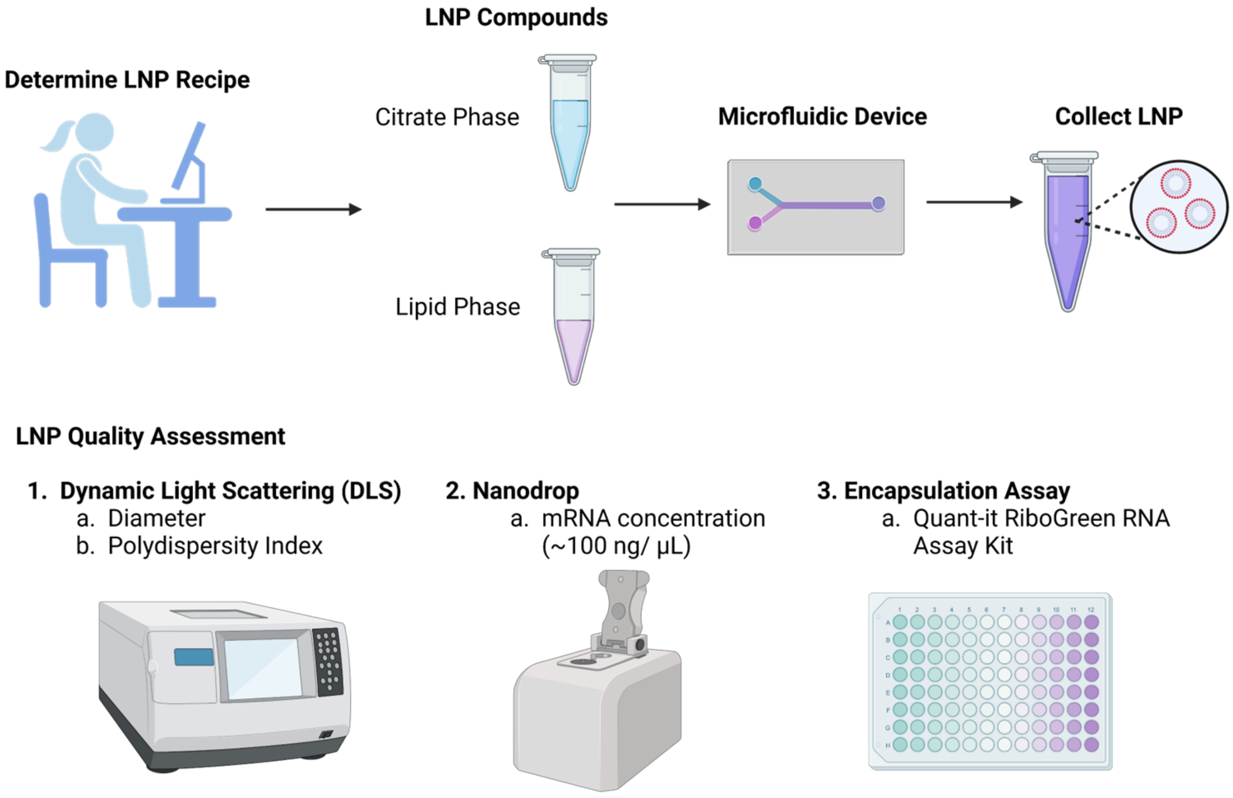


**Fig. S10. Steps for LNP formulation and quality assessment.** The LNP recipe was determined, and the corresponding citrate and lipid phases were combined in the microfluidic device for LNP formulation. The quality of the formulated LNP was evaluated using dynamic light scattering (DLS) to measure the diameter and polydispersity index (PDI), nanodrop to measure the mRNA concentration, and encapsulation assay to quantify the corresponding encapsulation efficiency.

**TABLES**

| LNP# | Lipomer | Cholesterol | PEG | Helper lipid | Lipomer mole % | Cholesterol mole % | PEG mole % | Helper lipid mole % |
|---|---|---|---|---|---|---|---|---|
| LNP1 | cKK-E12 | Cholesterol | C14PEG2K | DSPC | 30 | 30 | 2.5 | 37.5 |
| LNP2 | cKK-E12 | Cholesterol | C14PEG2K | DSPC | 35 | 18 | 2.5 | 44.5 |
| LNP3 | cKK-E12 | Cholesterol | C14PEG2K | DSPC | 45 | 42 | 2.5 | 10.5 |
| LNP4 | cKK-E12 | Cholesterol | C14PEG2K | DSPC | 50 | 35 | 2.5 | 12.5 |
| LNP7 | cKK-E12 | Cholesterol | C14PEG2K | DOPE | 30 | 30 | 2.5 | 37.5 |
| LNP11 | cKK-E12 | Cholesterol | C14PEG2K | DOPE | 52.5 | 15 | 2.5 | 30 |
| MC3 | MC3 | Cholesterol | C14PEG2K | DSPC | 50 | 38.5 | 1.5 | 10 |

**Table S1.** Compounds and molar ratios included in lead LEC-specific LNP candidates, namely LNP1, LNP2, LNP3, LNP4, LNP7, LNP11, MC3. Unique LNPs were formulated with different combinations of compounds and molar ratios of ionizable lipid, cholesterol, PEG, and helper lipid.

| Type of LN | Administration | LEC Count | aVHH$^+$ Cell Count |
|---|---|---|---|
| ALN | Saline | 2182 | 37 |
| | LNP1 | 1022 | 543 |
| | LNP2 | 1467 | 697 |
| | LNP3 | 948 | 134 |
| | LNP4 | 997 | 134 |
| | LNP7 | 2757 | 977 |
| | LNP11 | 372 | 37 |
| BLN | Saline | 2016 | 39 |
| | LNP1 | 2139 | 376 |
| | LNP2 | 2169 | 715 |
| | LNP3 | 1971 | 572 |
| | LNP4 | 3319 | 159 |
| | LNP7 | 2204 | 743 |
| | LNP11 | 3631 | 239 |
| PLN | Saline | 95 | 2 |
| | LNP1 | 388 | 139 |
| | LNP2 | 105 | 31 |
| | LNP3 | 771 | 144 |
| | LNP4 | 118 | 49 |
| | LNP7 | 782 | 292 |
| | LNP11 | 230 | 52 |

**Table S2.** Average LEC count and LEC/aVHH$^+$ cell count for lead LEC-specific LNP candidates, namely LNP1, LNP2, LNP3, LNP4, LNP7, and LNP11.

| | | Cell Type Count | | | | | | | | |
|---|---|---|---|---|---|---|---|---|---|---|
| Type of LN | Administration | Macrophages | Monocytes | cDC2 | cDC1 | DNs | FRCs | LECs | BECs | VECs |
| ALN | Saline | 4036 | 1294 | 2914 | 1008 | 114907 | 6175 | 7099 | 237 | 1269 |
| | Free aVHH | 4970 | 1606 | 4155 | 1356 | 270150 | 6045 | 10462 | 744 | 3796 |
| | MC3 | 4013 | 67256 | 4264 | 2682 | 155750 | 7848 | 7993 | 145 | 1995 |
| | LNP7 | 4665 | 10492 | 6018 | 3885 | 337655 | 8294 | 7802 | 167 | 1879 |
| BLN | Saline | 7113 | 2666 | 8368 | 2474 | 105104 | 13394 | 10007 | 296 | 2267 |
| | Free aVHH | 10295 | 4097 | 11666 | 3698 | 168662 | 15316 | 16065 | 574 | 6351 |
| | MC3 | 8562 | 6471 | 12017 | 4630 | 224232 | 21738 | 15810 | 249 | 5418 |
| | LNP7 | 7940 | 4129 | 12104 | 3390 | 227405 | 14637 | 10541 | 310 | 3433 |
| PLN Contralateral | Saline | 1790 | 728 | 1417 | 541 | 18325 | 2056 | 3034 | 3475 | 488 |
| | Free aVHH | 1069 | 439 | 915 | 636 | 15212 | 1273 | 3686 | 45 | 624 |
| | MC3 | 1302 | 651 | 1486 | 719 | 13422 | 2742 | 1267 | 28 | 381 |
| | LNP7 | 1784 | 927 | 1934 | 912 | 29328 | 2412 | 4061 | 62 | 1111 |
| PLN Injected | Saline | 2054 | 599 | 1330.25 | 336 | 28092 | 1791 | 3272 | 56.25 | 632 |
| | Free aVHH | 4425 | 1737 | 1680 | 874 | 27735 | 1990 | 4153 | 70.5 | 1002 |
| | MC3 | 7739 | 25134 | 2737 | 2445 | 34778 | 4618 | 4220 | 72 | 1091 |
| | LNP7 | 7568 | 25225 | 3605 | 2586 | 69388 | 5089 | 5042 | 85 | 1058 |
| PLV | Saline | NA | NA | NA | NA | NA | NA | 185 | NA | NA |
| | Free aVHH | NA | NA | NA | NA | NA | NA | 188 | NA | NA |
| | MC3 | NA | NA | NA | NA | NA | NA | 189 | NA | NA |
| | LNP7 | NA | NA | NA | NA | NA | NA | 269 | NA | NA |

**Table S3.** Average cell count for the various cell types found in ALN, BLN, PLN Contralateral (non-injected), PLN injected, and PLV for saline-, free aVHH-, MC3-, and LNP7-injected animals.

| | | aVHH$^+$ Cell Count | | | | | | | | |
|---|---|---|---|---|---|---|---|---|---|---|
| **Type of LN** | **Administration** | **Macrophages** | **Monocytes** | **cDC2** | **cDC1** | **DNs** | **FRCs** | **LECs** | **BECs** | **VECs** |
| **ALN** | **Saline** | 43 | 3 | 31 | 21 | 8.5 | 35 | 89 | 1 | 3 |
| | **Free aVHH** | 45 | 3 | 106 | 37 | 52 | 38 | 406 | 3 | 15 |
| | **MC3** | 320 | 876 | 437 | 31 | 31 | 44 | 263 | 1 | 18 |
| | **LNP7** | 201 | 1142 | 781 | 82 | 55 | 62 | 192 | 1 | 14 |
| **BLN** | **Saline** | 160 | 5 | 492 | 246 | 6 | 49 | 156 | 5 | 13 |
| | **Free aVHH** | 150 | 10 | 170 | 75 | 10 | 67 | 226 | 115 | 44 |
| | **MC3** | 287 | 20 | 162 | 47 | 14 | 65 | 254 | 3 | 41 |
| | **LNP7** | 152 | 28 | 578 | 441 | 14 | 57 | 409 | 7 | 21 |
| **PLN Contralateral** | **Saline** | 50 | 2.5 | 37 | 52 | 3 | 17 | 65 | 0.5 | 2 |
| | **Free aVHH** | 37 | 2 | 97 | 149 | 5 | 9 | 158 | 1 | 12 |
| | **MC3** | 25 | 4 | 43 | 35 | 12 | 68 | 45 | 1 | 4 |
| | **LNP7** | 28 | 2 | 82 | 75 | 8 | 16 | 89 | 2 | 8 |
| **PLN Injected** | **Saline** | 73 | 2 | 31 | 13 | 6 | 21 | 98 | 1 | 7 |
| | **Free aVHH** | 78 | 11 | 29 | 11 | 10 | 21 | 79 | 1 | 8 |
| | **MC3** | 772 | 2755 | 521 | 86 | 78 | 296 | 591 | 1 | 20 |
| | **LNP7** | 888 | 3681 | 1677 | 978 | 202 | 568 | 1690 | 3 | 77 |
| **PLV** | **Saline** | NA | NA | NA | NA | NA | NA | 3 | NA | NA |
| | **Free aVHH** | NA | NA | NA | NA | NA | NA | 3 | NA | NA |
| | **MC3** | NA | NA | NA | NA | NA | NA | 18 | NA | NA |
| | **LNP7** | NA | NA | NA | NA | NA | NA | 104 | NA | NA |

**Table S4.** Average aVHH$^+$ cell count for the various cell types found in ALN, BLN, PLN Contralateral (non-injected), PLN injected, and PLV for saline-, free aVHH, MC3-, and LNP7-injected animals.
